# Interpretable biophysical neural networks of transcriptional activation domains separate roles of protein abundance and coactivator binding

**DOI:** 10.1101/2025.09.19.677413

**Authors:** Claire LeBlanc, Pooja Agarwal, Jack Demaray, Gean Hu, Marissa Zintel, Angelica Lam, Joel Enrique Castro Hernandez, Max Staller

**Author notes:** University of California San Francisco, San Francisco, CA 94158.

## Abstract

Deep neural networks have improved the accuracy of many difficult prediction tasks in biology, but it remains challenging to interpret these networks and learn molecular mechanisms. Here, we address the interpretability challenges associated with predicting transcriptional activation domains from protein sequence. Activation domains, regions within transcription factors that drive gene expression, were traditionally difficult to predict due to their sequence diversity and poor conservation. Multiple deep neural networks can now accurately predict activation domains, but these predictors are difficult to interpret. With the goal of interpretability, we designed simple neural networks that incorporated biophysical models of activation domains. The simplicity of these neural networks allowed us to visualize their parameters and directly interpret what the networks learned. The biophysical neural networks revealed two new ways that arrangement (i.e. the sequence grammar) of activation domain controlled function: 1) hydrophobic residues both increase activation domain strength and decrease protein abundance, and 2) acidic residues control both activation domain strength and protein abundance. Notably, the biophysical neural networks helped us to recognize the same signatures in complex interpreters of the deeper neural networks. We demonstrate how combining biophysical and deep neural networks maximizes both prediction accuracy and interpretability to yield insights into biological mechanisms.

## Introduction

Deep neural networks (NNs) have transformed molecular biology, with applications varying from protein structure prediction^1–3^ to single-cell transcriptomics^4,5^. Deep NNs learn complex patterns using millions of parameters, but this complexity also makes it difficult to interpret how and why they are making specific predictions^6,7^. The lack of interpretability surrounding these NNs is of particular concern in biological research, where one of the fundamental goals is not just to make accurate predictions, but also mechanistic understanding of the underlying processes^5^.

The field of machine learning interpretability has emerged to shed light on what these NNs are learning^6–8^. There are two common approaches for interpreting biological NNs: either applying post-hoc interpretability methods^8,9^ or building biological knowledge into the NN architecture^10–13^. In this work, we demonstrate that both simple NNs and NNs that incorporate biological models enhance interpretability and our understanding of the underlying biology.

We focused on the biological challenge of predicting activation domains within transcription factors (TFs) from protein sequence. Activation domains are located within intrinsically disordered regions of TFs and interact with coactivator proteins to turn on transcription of a gene^14,15^. Traditionally, the problem of identifying activation domains from sequence has been difficult because they are disordered, diverse, and poorly conserved^16,17^. High-throughput screens have uncovered many sequence features of acidic activation domains, including a balance between acidic and aromatic amino acids^14,18–27^. The large amount of data generated from these screens has also been used to train deep NNs that accurately predict activation domains from protein sequence^21,25,28,29^.

We sought to interpret NNs to learn the sequence grammar controlling activation domain function. Sequence grammar is the specific arrangements of amino acids that lead to function^30,31^. We do not yet fully understand the grammar of activation domains^14^. Although amino acid composition is important for function, grammar is also important in IDRs^31,32^ and activation domains^14,33^. Sequences containing acidic and aromatic amino acids are not always activation domains^23,25,26^, and predicting activation domains using amino acid composition alone does not achieve the same accuracy as NNs^33^. The high performance of NNs compared to composition-based predictors suggests that they are learning some grammar. While previous NNs for activation domain prediction have been interpreted using post-hoc methods, these interpretations had limited utility and did not reveal a strong grammar signal^21,28^. We hypothesize that building biological knowledge into NNs will improve interpretability and reveal the grammar of activation domains.

In this work, we used interpretable NNs to test our understanding of the features that control activation domain function. Our goal was not to outperform the existing predictors, but merely to increase interpretability while achieving similar accuracy. We tried two approaches: 1) designing the simplest possible NN that maintained prediction accuracy, and 2) incorporating biophysical models into the simple NN. The simple NNs have minimal parameters that can be directly examined. By directly interpreting these parameters, we confirm known features of activation domains, such as the importance of acidic and aromatic amino acids. The biophysical NNs additionally reveal two grammar signals in activation domains: negative amino acids have position-specific effects on TF abundance and coactivator binding, and hydrophobic amino acids have conflicting C-terminal effects on TF abundance and coactivatior binding. Finally, these BiophysicalNNs help us to make sense of the post-hoc interpretations of deeper, more accurate NNs. Our results highlight the utility of simple, interpretable NNs for learning molecular mechanisms that govern protein function.

## Results

### The data and neural network architecture

We trained our NN using data from a previous screen measuring two dimensions of activation domain function: TF **abundance** and activation domain **strength**^17^. We define strength as the amount of reporter gene expression in our screen. In this screen, *S. cerevisiae* cells were transformed with synthetic TFs that contain an mCherry to quantify TF abundance, a DNA-binding domain targeted to a synthetic reporter locus coding for GFP, an estradiol-binding domain for inducible activation, and a variable C-terminal 40-amino acid sequence that may act as an activation domain (**Figure 1A**). Two Sort-seq^23^ experiments were performed to measure the amount of mCherry and GFP expression for each variable 40-amino acid sequence (**Methods**). Our NNs take as input the variable 40-amino acid sequence and predict either 1) the amount of GFP expression, which corresponds to the activation domain strength, 2) the amount of mCherry expression, which corresponds to the overall TF abundance, or 3) both of these quantities. For all the NN variations, we used the same basic architecture building block (**Figure 1B**).

**Figure 1:**
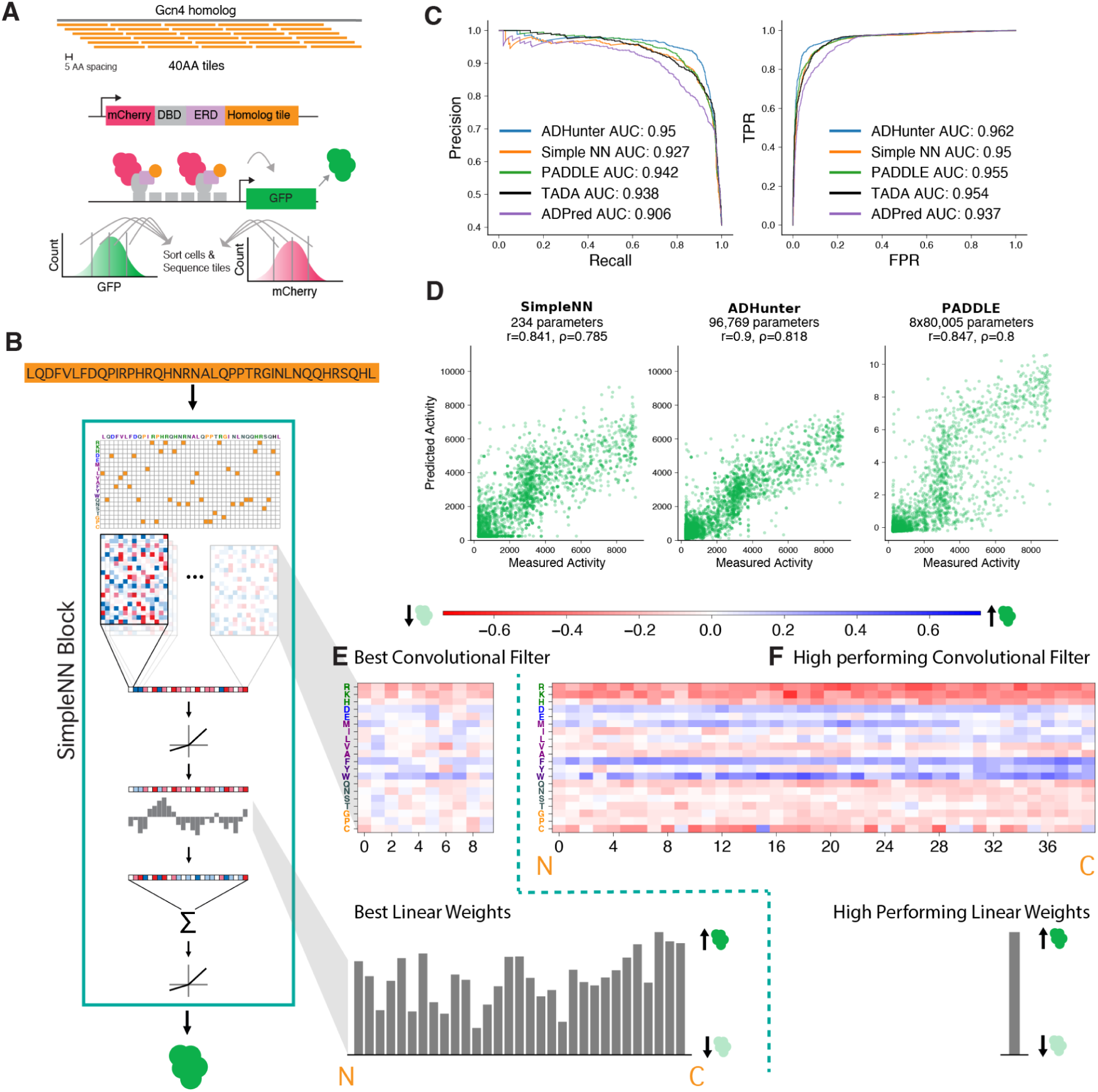
The SimpleNN accurately predicts activation domain strength and learns known features of activation domains. **A)** The experimental screen used to generate the training data. Cells are transformed with synthetic transcription factors containing an mCherry fluorophore, a DBD targeting a GFP reporter, an estrogen receptor domain, and a 40aa tile. Cells are sorted based on GFP or mCherry and then sequenced. **B)** The SimpleNN block architecture used to predict GFP expression for a given 40aa tile. The same block is used in all other NNs. **C)** Precision-recall curve and ROC curve for the SimpleNN and published deep NNs on an unseen test set. **D)** Scatter plots comparing the SimpleNN, ADhunter-GFP and PADDLE on the test set. **E)** SimpleNN convolutional filter weights and dense layer weights for the best performing NN. N and C mark which weights were applied to the features corresponding to the N and C terminus of the original sequence. Colorbar shows the convolutional filter scale, with red indicating negative weights (leading to a smaller prediction) and blue indicating positive weights (leading to a larger prediction). For this smaller convolutional filter, the linear weights capture most of the position-specific information. **F)** Convolutional filter weights and linear layer weights for another NN with a filter size of 40aa. Colorbar is the same as for **1E**. Here, the convolutional filter captures all position information. N and C markers indicate the N and C terminus of the original sequence.

This block takes as input the one-hot encoded activation domain sequence and applies a convolutional filter, followed by a parametric ReLU activation function, a dense layer with a single output, and a final parametric ReLU activation function. This architecture is used in all of our NNs to predict either the total TF abundance (mCherry fluorescence), the activation domain strength (GFP fluorescence), or an intermediate biophysical value (the equilibrium constant between states). We chose to use a single convolutional filter because a single dense layer could not as accurately predict the phenotype, and multiple convolutional filters rapidly became uninterpretable. Additionally, in NNs of DNA sequence, convolutional filters excel at detecting TF binding motifs^13^. We reasoned that if protein motifs play a large role in this dataset, we would detect them. We systematically varied the size of the convolutional filter and the initialization conditions (see the **Methods**). The random initialization of the NN had a larger impact on performance, and most convolutional filter sizes fell within the noise caused by random initialization (**Figure S1**).

### The simple neural network for predicting activation domain strength

First, we trained a NN to predict activation domain strength (the GFP expression) using a single building block (**Figure 1B**). We started with this simple NN (SimpleNN) as a baseline, so that we could evaluate whether the biophysical NNs provided any additional benefit. Although our main goal was to build an interpretable NN, we benchmarked performance of the SimpleNN against existing predictors. We evaluated our best performing SimpleNN on an unseen test dataset. Our SimpleNN is a slight improvement over linear regression on the one-hot encoding (Pearson correlation of 0.841 vs 0.801), likely because of the non-linearities captured by NNs.

We also compared it with three published deep NNs: TADA^28,29^, PADDLE^25^, and ADPred^21^, and one in-house predictor trained on the same dataset: ADhunter-GFP (**Methods**). ADhunter-GFP, the SimpleNN, and PADDLE were trained as regressors while TADA and ADPred were trained as classifiers. We compared the ability of our NN to make quantitative predictions (ADhunter-GFP, PADDLE) and to classify sequences as activation domains (ADhunter-GFP, TADA, PADDLE, and ADPred). The best performing SimpleNN performs slightly worse than ADhunter-GFP and PADDLE on both regression and classification on the test set (**Figure 1C, D**) while using 300x and ∼2500x fewer parameters, respectively (234 for the SimpleNN vs. 96,769 ADhunter and 8 ensembled NNs of 80,005 for PADDLE). The SimpleNN also performs comparably to the other predictors on the classification test (**Figure 1C**). Like the deep NNs, the SimpleNN generalizes to new datasets^27,28^ with slightly worse performance (**Table S1**).

Next, we analyzed the parameters of the SimpleNN to determine what the SimpleNN is learning (**Figure 1E, F**). The simplicity of the network, two layers, allows direct visualization of the NN parameters. For a detailed description of how these parameters function in the context of the NN, see **Methods**. Briefly, the convolutional filter identifies important amino acid features. The parameters of the dense layer weigh these features based on where they appear in the input 40 amino acid (aa) sequence. As the convolutional filter size increases, the convolutional filter also begins to pick up positional effects. At the largest convolutional filter size (40aa), the convolutional layer is mathematically equivalent to a dense layer with a single output node.

The convolutional filter weights and dense layer weights are shown for two high-performing NNs with different convolutional filter sizes in **Figure 1E** and **1F**. When looking at these parameters, we recover known features of acidic activation domains (**Figure 1E, F**).

Both NN convolutional filters learn the detrimental effect of positive amino acids as previously described^22,27,34^. Additionally, both filters learn that phenylalanine, tryptophan, and aspartic acid increase activation domain strength. These signals align with previous observations that acidic and aromatic amino acids are essential for activation domain function^14,18–27^. These same patterns were observed across NNs with various random initializations and filter sizes (**Figure S2**). The convolutional filters are not picking up on specific protein motifs, and instead are picking up composition-based patterns. When we looked at the linear weights, features at the end of the sequence had a slightly greater impact on activation domain strength, as has been previously observed^27^ (**Figure 1E, S2**). These results indicate that for the task of activation domain prediction, a two-layer NN is interpretable, captures known features of activation domains, and has similar accuracy as more complex NNs.

### The simple neural network for predicting transcription factor abundance

Next, we built simple NNs for the second phenotype in the screen: transcription factor abundance (**Figure 1A**). To identify the sequence features that determine transcription factor abundance, we retrained our SimpleNN to predict the TF abundance (SimpleNN-abund, **Figure 2A**). The abundance data was much noisier than the GFP data (replicate Pearson correlation of 0.568), and our best performing NN had a Pearson correlation of 0.532 on unseen test data (**Figure 2B**). For a comparison to a deep NN, we retrained ADhunter to predict the mCherry value (ADhunter-Cherry). ADhunter-Cherry once again achieved slightly higher performance while using almost 300x more parameters (**Figure 2B**).

**Figure 2:**
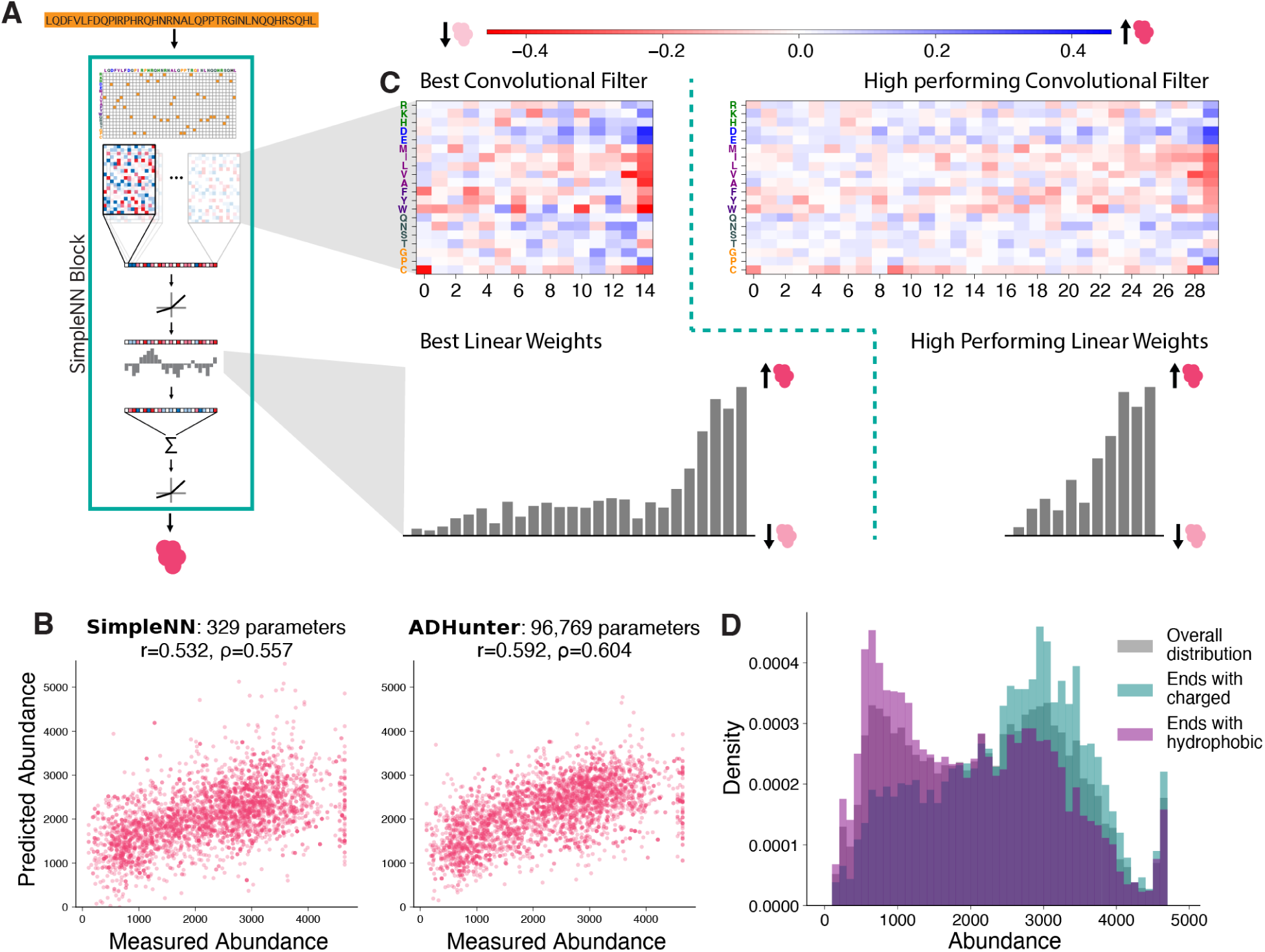
The SimpleNN for abundance reveals influence of activation domain sequence on TF abundance. **A)** The same SimpleNN block architecture as used in Figure 1 is used to predict mCherry expression for a given 40aa tile. **B)** Scatter plots comparing the SimpleNN for abundance and ADhunter-Cherry on a held out test set. **C)** Neural network parameters for two high performing SimpleNNs with different convolutional filter sizes. Colorbar shows the convolutional filter scale, with red indicating negative weights (leading to a smaller prediction) and blue indicating positive weights (leading to a larger prediction). **D)** Abundance distributions of synthetic TFs that end with a charged amino acid versus a hydrophobic amino acid. Hydrophobic amino acids are M, I, L, V, A, F, Y and W. Charged amino acids are R, K, D, and E.

The interpretability of the SimpleNN-abund led us to find that the last few amino acids of the sequence had the largest impact on TF abundance. As seen in the NN parameters, the presence of hydrophobic amino acids in the last ∼3 positions leads to a predicted decrease in abundance, while charged amino acids lead to a predicted increase in abundance (**Figure 2C, S3**). When we went back to our experimental data, we found that sequences that ended in a hydrophobic amino acid had lower abundance than those ending in a charged amino acid, confirming that the NN had learned a trend in the data that we had previously missed (**Figure 2D**, **S4**). As the sequences acquired more C-terminal hydrophobic amino acids, they became even less abundant (**Figure S4**). This trend was distinct from the effect of hydrophobic amino acids throughout the sequence, which also decreased transcription factor abundance to a lesser extent (**Figure S4**). By interpreting our simple NN, we were able to uncover strong biological signals that we had previously missed*.

Because all synthetic TFs are under control of the same promoter, differences in abundance are most likely due to protein destabilization and degradation. Supporting this hypothesis, a recent paper performing a high-throughput screen of transcription factor abundance found that treatment with a proteosome inhibitor increased the abundance of almost all TFs, indicating that transcription factor abundance in these screens is frequently controlled through protein degradation^35^. We analyzed the abundance data from two screens in human cells from the same group^35,36^ and found that sequences ending in hydrophobic amino acids are enriched in low abundance sequences, consistent with the patterns in our data (**Figure S5**). We also found that C-terminal *S. cerevisiae* activation domains were enriched for acidic residues in the last ten positions (*p*-value=0.015, Mann–Whitney U test, one-tailed, **Figure S6**). This difference in terminal negative charge suggests that negative amino acids in C-terminal activation domains could also play a role in preventing TF degradation.

### Incorporating Biophysical Models into Neural Network Architecture

To further increase interpretability, we next designed NNs that incorporated a biophysical model of activation domains (the BiophysicalNNs). Unlike with the SimpleNNs, where these values were predicted by two separate NNs, these biophysical NNs simultaneously predict TF abundance and activation domain strength. We tested two biophysical models, both inspired by our acidic exposure model of activation domain function^26^. In the acidic exposure model, key aromatic and leucine residues make contacts with hydrophobic surfaces of coactivators in the bound state. However, in the unbound state, these same hydrophobic residues interact with each other to drive a collapsed, inactive state. The acidic residues counteract the collapse with electrostatic repulse and favorable free energy of solvation, promoting an exposed state where the hydrophobic residues are available to bind the coactivator^26^. We formalized the acidic exposure model as a three-state biophysical model (**Figure 4A**). As an intermediate, we first studied a two-state binding model where the activation domains exist in two states: unbound or bound to coactivator (**Figure 3A**). Only activation domains in the second state (bound to coactivator) are able to activate GFP expression^37–39^.

**Figure 3:**
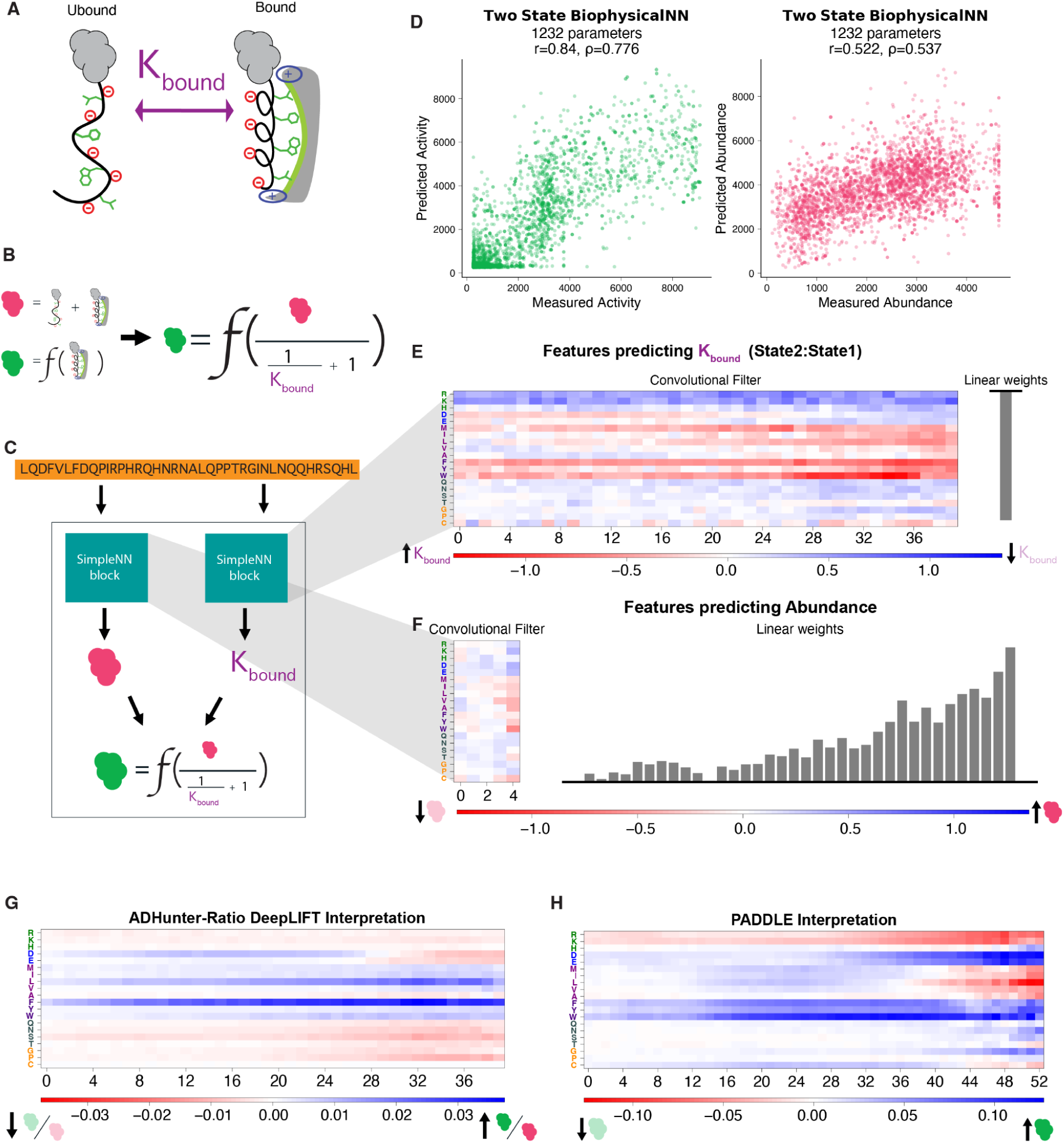
The two-state model of transcription factor activation domains learns amino acid tradeoffs. **A)** The two-state biophysical model of transcription factor activation domains. **B)** Assumptions used to derive the relationship between the biophysical model, the mCherry TF abundance, and the GFP activation. **C)** The architecture of the two-state BiophysicalNN. The SimpleNN block is shown in Figure 1 and 2. We tried multiple functions for *f* and found that a hill function led to the highest accuracy. **D)** Scatterplots showing the predictive performance of the best performing two-state BiophysicalNN. Activation domain strength and abundance are predicted simultaneously. **E)** The trainable parameters predicting the equilibrium constant between the two states of the biophysical model for the best performing two-state BiophysicalNN. A convolutional filter size of 40aa performed best across random seeds. Note that the linear weight is negative, reversing the effect of the weights in the convolutional filter. **F)** Parameters from the same best performing two-state BiophysicalNN that predict the abundance. **G)** Average of ADhunter-ratio DeepLIFT scores for all activating tiles. **H)** Post-hoc interpretation of PADDLE. Each amino acid in five transcription factors was mutated to every other amino acid and the average effect on the final prediction was calculated.

**Figure 4:**
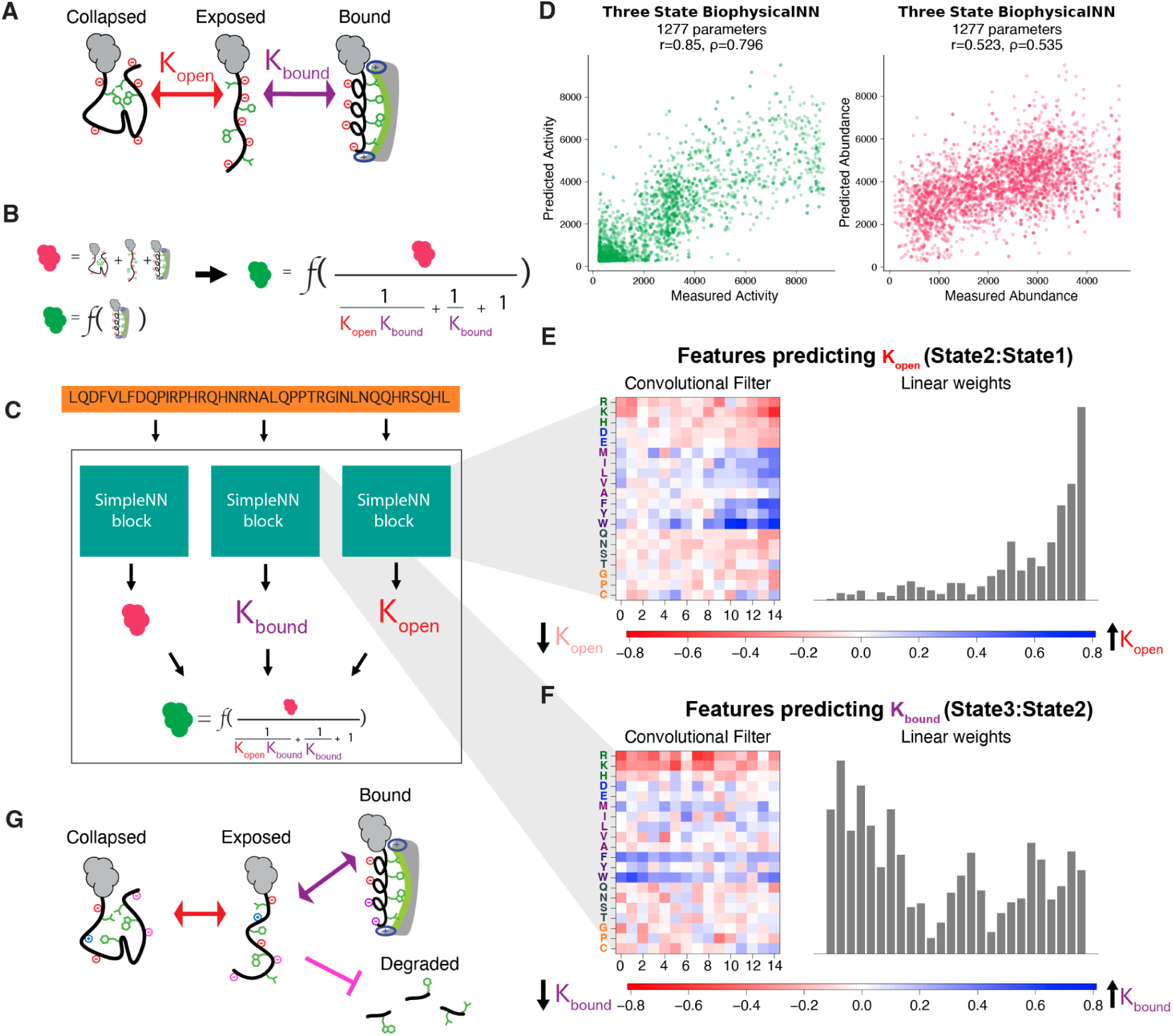
The three-state BiophysicalNN does not learn the acidic exposure model. **A)** The three-state acidic exposure model of activation domains. **B)** Assumptions used to derive the relationship between the acidic exposure model, the TF abundance, and the GFP activation. **C)** The architecture of the three-state BiophysicalNN. The SimpleNN block is shown in Figures 1 and 2. We tried multiple functions for *f* and found that a hill function led to the highest Pearson correlation. **D)** Scatterplots showing the predictive performance of the best performing three-state BiophysicalNN. activation domain strength and abundance are predicted simultaneously. **E)** The trainable parameters predicting the equilibrium constant between state 1 and state 2 (collapsed and expanded) of the best performing three-state Biophysical-NN, which used a filter size of 15. **F)** The trainable parameters for the same best performing NN predicting the equilibrium constant between state 2 and state 3 (expanded and bound). **G)** An updated biophysical model where negative amino acids both prevent hydrophobic collapse and promote TF abundance.

We derived an equation relating the equilibrium constants between the states of the biophysical models to the experimental measures of activation domain strength and transcription factor abundance (**Figure 3B, 4B, S7**). We used the equilibrium constants because we wanted the intermediate biophysical values to be independent of overall abundance. The BiophysicalNNs predict protein abundance and the equilibrium constants using the basic building blocks described earlier. The predicted equilibrium constants are used to calculate the amount of transcription factor in the bound state. Finally, a hill function is applied to predict the total amount of GFP expression from the amount of bound TF (See **Methods**). The architectures for the two-state and three-state BiophysicalNN are shown in **Figure 3C** and **Figure 4C**, respectively.

### Biophysical Neural Networks retain prediction accuracy

The best performing BiophysicalNN containing the two-state activation domain model had similar accuracy compared to the SimpleNN (**Figure 3D**). For larger filter sizes, the two NNs have very similar accuracies (**Figure S8**). The requirement for larger filter sizes in the two-state BiophysicalNN is likely due to the position-specific grammar signals detected by the two-state BiophysicalNN, which can only be captured by larger convolutional filters that incorporate positional information. These grammar signals will be discussed in the following section.

The three-state BiophysicalNN also performed comparably to the SimpleNN and the best performing two-state BiophysicalNN (**Figure 4D, S8**). Like the SimpleNN, both BiophysicalNNs performed similarly to the deeper NNs with many fewer parameters. Both the BiophysicalNNs generalized slightly better than the SimpleNN on new data (**Table S1**). Overall, all NN variations performed similarly, with individual differences likely coming from differences in random initialization rather than the inherent benefits of one architecture over the other.

### Biophysical Neural Networks reveal activation domain grammar

Next, we investigated what our BiophysicalNNs were learning, starting with the two-state BiophysicalNN. We specifically focused on the sequence features that promoted a higher equilibrium constant between the bound and unbound states by looking at the convolution filter and dense layer weights (**Figure 3E**). Importantly, the dense layer has a negative weight, so the effect of the convolutional filter weights on the final prediction is reversed.

The two-state BiophysicalNN learned that tryptophan and phenylalanine residues throughout the sequence are all important in promoting coactivator binding, similar to the GFP SimpleNN (**Figure 3E**). Also, like the SimpleNN, positive residues throughout the sequence decrease the amount of TF in the bound state.

Unlike the SimpleNN, the BiophysicalNN learned that negative amino acids have two conflicting functions with position-specific dependence. Negative amino acids at the beginning of the sequence are important for binding to coactivator, while negatives at the end of the sequence are important for promoting TF abundance (**Figure 3E, F-rows for D and E**). We also averaged parameters across NNs with different random initializations and found the same trends (**Figure S9**). In contrast, the SimpleNN learned that negative amino acids throughout the sequence increase activation domain strength (**Figure 1E, F**). For negative amino acids, the GFP SimpleNN convolutional filter resembles a combination of the abundance and equilibrium constant convolutional filters in the two-state BiophysicalNN (**Figure 3E, F**). The two-state BiophysicalNN can separate the two functions of acidic residues.

Beyond the grammar of negatives, we also found other differences between the SimpleNN and the two-state BiophysicalNN. The BiophysicalNN learns that methionine throughout the sequence promotes coactivator binding, while isoleucine, leucine, and valine at the end of the sequence promote coactivator binding (**Figure 3E**). The SimpleNN finds that valine prevents activation, likely because although valines at the end of the sequence promote binding, those same valines also reduce TF abundance (**Figure 1E, F**). By separating the contributions to coactivator binding and abundance, the BiophysicalNN suggests a dual role for valine.

We were curious whether the predicted equilibrium constant was biologically meaningful. In the absence of experimental data on the equilibrium constants for our tiles, we instead used FINCHES^40^ to predict the strength of the interaction between the 40 amino acid tile and the Med15 coactivator ABD1. FINCHES predicts the interactions between IDRs and partner proteins using molecular dynamics force fields, and was shown to accurately predict IDR-mediated interactions^40^. We found that the equilibrium constant is correlated with the FINCHES predicted interaction strength (**Figure S10**). In the absence of experiments, we caution against interpreting the equilibrium constants as a true biophysical measure, but the relationship between the interaction strength and the equilibrium constant of binding is intriguing.

### Biophysical Neural Networks help to interpret more complicated NNs

The BiophysicalNNs helped us to understand the post-hoc interpretations of the deeper NNs, revealing that the deep NNs also learned bidirectional amino acid contributions. We performed a DeepLIFT^9^ interpretation of ADhunter-ratio (**Methods**), our in-house predictor trained on data from cells sorted on the GFP:mCherry ratio (the GFP expression normalized by the mCherry signal). Ratio sorting is our most commonly collected and highest resolution data, and ADhunter-ratio is our workhorse predictor. Note that ADhunter-ratio is different than the newly released ensemble version of ADhunter^41^ (ADhunter-ensemble), which was trained on a different dataset. DeepLIFT scores calculate how much each amino acid in the input is contributing to the final prediction^9^. Averaging all DeepLIFT scores for the Gcn4 dataset indicated that ADhunter-ratio is learning a bifunctional impact of negative amino acids (**Figure 3G**). This result was surprising, because previous analyses had suggested that negatives throughout the sequence increase activation domain strength^18–27^. This trend aligns, however, with our BiophysicalNN interpretation. Negative amino acids at the C-terminus increase the total abundance of the transcription factor, counterintuitively leading to a decrease in the GFP to mCherry expression ratio. The ADhunter-ratio DeepLIFT also finds that aromatic and bulky hydrophobic amino acids have the strongest effect when located at the end of the sequence.

The BiophysicalNN also helped us to understand these trends more deeply: C-terminal hydrophobic amino acids both decrease abundance and increase coactivator binding. Finally, the trends in our interpretation of ADhunter-ratio match the trends in the interpretation of ADhunter-ensemble^41^, suggesting these patterns may result from ratio sorting, and not the data itself. The BiophysicalNN identified two instances of bidirectional amino acid contributions in ADhunter-ratio, interpreting the interpreters.

We also performed an interpretation of another deep NN trained to predict activation domains, PADDLE^25^. We took five known activation domains, ran PADDLE on all possible single amino acid mutations, and averaged the effect of each perturbation on the output (See **Methods** for details). We chose this approach because PADDLE was not amenable to DeepLIFT because the original code used to write and train the NN was not available. Additionally, this approach, applied to ADhunter-ratio, captures similar (although not identical) trends to the DeepLIFT interpretation (**Figure S11**). Based on this PADDLE interpretation, C-terminal non-aromatic hydrophobic amino acids tend to decrease the predicted activation domain strength (**Figure 3H**). Initially, this trend was counterintuitive, as previous analyses have found that bulky hydrophobic amino acids positively contribute to activation domain function^17,19,21–23,25–28^. From our BiophysicalNN, however, we propose that these hydrophobic residues may decrease transcription factor abundance, leading to lower gene expression. In the training data for PADDLE, Sanborn et al.^25^ attempted to control for transcription factor abundance by using an mCherry gating scheme, but based on our interpretation of PADDLE, we hypothesize that there may have still been variability in TF abundance that contributed to reporter gene expression. These two examples demonstrate how BiophysicalNNs can be combined with more sophisticated post-hoc interpretation techniques to enhance our understanding of what deeper NNs are learning.

### The Three-State Biophysical NN is not learning the acidic exposure model

We set up the three-state BiophysicalNN to formalize the acidic exposure model (**Figure 4A**). In this biophysical model, negative amino acids promote the exposed state (state 2) and aromatic amino acids promote the bound state (state 3) and the collapsed state (state 1). We did not enforce this assumption in the NN, however, instead merely designing a NN with three states where the final state is the only state capable of activating gene expression (**Figure 4B, Figure S7**). When we investigated the parameters of the three-state BiophysicalNN, it was not learning the acidic exposure model (**Figure 4E, F**). We did not see a consistent pattern of negative amino acids promoting the exposed state and aromatic amino acids promoting the bound state (**Figure S13**). Instead, the three-state BiophysicalNN seemed to be picking up on the position-specific effects of negative amino acids that were observed in the two-state BiophysicalNNs (**Figure S13**). Specifically, one filter learned that the negatives at the beginning of the sequence promote coactivator binding (**Figure 4F**) while the other learned that negatives at the end do not (**Figure 4E**).

There are many possible reasons that our three-state BiophysicalNN is not learning the acidic exposure model. Potentially, the BiophysicalNN is learning a different three-state model of transcriptional activation domains in which the final state promotes gene expression. For example, it could be learning a two step binding mechanism, where binding is initiated through electrostatic interactions^42^ then followed by hydrophobic interactions^37^. Alternatively, if the transitions between the exposed and collapsed states are rapid^43^, then there is a separation of time scales and the system is better approximated by a two-state model. Estimates of intramolecular reorganization rates in IDRs are faster than the diffusion limit, which many IDR binding interactions approach,^44^ so this separation of time scales is plausible.

The most likely explanation, however, is that our NN is not learning any three-state biophysical model. NNs are known to build in redundancy, where the same features are picked up by various parts of the network^45^. This tendency may explain why both of the filters are picking up on the importance of aromatic amino acids, as they are essential for activation. More innovative NN design, more complex loss functions, or more experimental measurements of intermediate states may be required to reduce redundancy and learn biophysical parameters.

## Discussion

As neural networks proliferate in biology and interpretability becomes more important, most interpretation approaches are either more sophisticated algorithms, or simple models that are designed for direct interpretation. Here, we designed directly interpretable simple and biophysical NNs that accurately predict transcriptional activation domains from protein sequence. These interpretable NNs revealed dual roles for hydrophobic and acidic amino acids. These dual roles were also present, but not previously appreciated, in the sophisticated interpretations of large NNs. In this way, the interpretable NNs helped us to understand the interpretations of large models.

We designed our NNs to predict two key functions of activation domains, protein abundance and activation domain strength. The SimpleNNs consist of a single convolutional and dense layer to predict either activation domain strength or TF abundance. The BiophysicalNNs, containing either a two or three-state model of activation domain function, predict both activation domain strength and TF abundance simultaneously. These NNs achieve similar accuracy to deep NNs with orders of magnitude fewer parameters. The SimpleNNs to predict activation domain strength learn known features of activation domains^14,18–27^ (such as the importance of acidic and aromatic amino acids), while the SimpleNNs to predict TF abundance and the BiophysicalNNs allow us to identify new signals in our data.

Previously, the impact of transcription factor abundance in high-throughput activation domain screens has either been ignored^21,22^ or been normalized out through ratio sorting^23,26,28^ or controlled for with gating^25^. Here, we took an alternative approach and used BiophysicalNNs to directly model the relationship between abundance and activation domain strength, which allows us to deconvolve bifunctional roles of amino acids and enhance our understanding of activation domain function (**Figure 4G**). For example, we found that hydrophobic amino acids, especially in the final C-terminal amino acids, both increase coactivator binding and decrease TF abundance. This finding aligns with previous studies that have found that activation domains tend to overlap with degrons^35,46,47^ and that high C-terminal hydrophobic content often decreases protein abundance^48,49^. These contradictory functions could be because C-terminal hydrophobic residues target proteins to the proteosome, or because the act of activating genes (via interactions with coactivators) leads to TF degradation, as previously suggested^50,51^.

Through our BiophysicalNN, we also identified a new grammar signal for negative amino acids. Negative amino acids at the N-terminal end of the activation domain increase AD-coactivator binding, while negative amino acids at the C-terminal end of the activation domain decrease AD-coactivator binding. The detrimental effects of C-terminal negative amino acids on AD-coactivator binding, however, are counteracted because C-terminal negative amino acids increase TF abundance. Two recent high-throughput screens of protein abundance in humans also found that negatively charged amino acids promote higher abundance and that acidic residues can counter the degron effect of hydrophobic amino acids in activation domains^35,36^. We propose that charge at the C-terminus of activation domains is more important for protein abundance than for binding to coactivator.

The bifunctional role of amino acids that we observed also allows us to reinterpret previous studies. For example, a deep mutational scan study of the CRX transcription factor found that replacing the C-terminal amino acids (WKFQIL) with aliphatic or negative amino acids increased activation domain strength^52^, despite the earlier observation that this motif is important for function^53^. This effect could be because of the balance between acidic and hydrophobic residues, as the authors suggested. Alternatively, we suggest that removing aromatic residues increases TF abundance, which subsequently increases activation domain strength.

Protein abundance is a necessary precondition for TF function. Many clinical mutations destabilize proteins, prompting efforts to improve the diagnosis of Variants of Uncertain Significance by combining deep mutational scans with measures of protein abundance^54^.

Clearly, TF abundance matters for activation, which is why we and others have tried to control for it with ratio sorting or gating narrowly on TF abundance. Depending on the normalization, each deep NN is learning a different abundance signature, indicating that abundance is important.

As has been previously noted, increased interpretability often leads to a decrease in accuracy^6^, and this work is no exception. Despite the mild decrease in predictive performance, we demonstrate that there is value in building interpretability and biophysical models into NNs and provide one framework for doing so. Additionally, we highlight the value of interpreting many NNs in parallel to ensure robust conclusions, as previously demonstrated^12,55,56^. We trained over 1800 NNs with varied architecture and random initializations. By interpreting all of these NNs in parallel, we were able to distinguish robust trends from noise.

This study complements a growing body of research on NN interpretability. Much research focuses on post-hoc interpretability methods, which can provide useful biological insight. For example, a DeepLIFT/DeepSHAP analysis of NNs trained to predict ATACSeq or DNASeq data revealed an enzyme bias in the data and enabled the authors to design a NN to correct for this bias^57^. For activation domains, the deep NN TADA was interpreted using SHAP, which revealed subclasses of activation domains and supported the acidic exposure model^28^.

Other research focuses on incorporating biological knowledge into NNs to increase interpretability. For example, Schmitt et. al designed a NN that incorporated a biophysical model, which provided insight into the spatial protein distributions affecting cell forces^56^. Many other biological NNs focus on building biological networks, such as networks of gene or protein interactions, into the architecture of NNs, which has also led to biological insight through post-hoc interpretation methods^55^. For example, DeepLIFT interpretation of a NN trained to predict prostate cancer disease state, P-NET, revealed both known and novel prostate cancer driver genes^58^. Post-hoc interpretations also have drawbacks, however, and often cannot distinguish biases in the network from real biology^12,59^. For example, some of the post-hoc interpretations from P-NET were shown to be artifactual^12^. Although techniques have been developed to mitigate artifacts^12,59^, these challenges highlight the value of directly interpreting NN parameters.

This work is most similar to (and inspired by) the recent development of biophysical NNs used to predict genotype-phenotype maps^10,11^. These biophysical NNs, MoCHI^11^ and MAVE-NN^10^, are trained on massively parallel reporter assay data. To map sequence to a biophysical value, MAVE-NN and MoCHI predict intermediate latent phenotypes or biophysical values (such as Gibbs free energy) linearly (i.e. adding the effects of single and double mutations). These NNs are easily interpretable, but require data for all single and many double mutants in reference to a single WT sequence. In contrast, we permit more complexity in the prediction of the biophysical value, while still maintaining enough simplicity to directly interrogate our network. We introduced more complexity because we found a simple linear relationship did not fit our data as well as the non-linear convolutional and dense layers, likely because our sequences were too diverse for the simple additive assumption. Another similar biophysical NN, BoltzNet, also uses a single convolutional filter to predict an intermediate biophysical value, in this case, the equilibrium constant of a TF binding to DNA^13^. They showed that their NN correctly learns biophysical parameters and like us, they directly interpreted the convolutional filter weights^13^. In our BiophysicalNNs, we incorporate features of both BoltzNet^13^, by using a convolutional filter to predict an equilibrium constant, and of MoCHI^11^, by having our BiophysicalNNs simultaneously predict two measures of activation domain function. Combining these approaches is what allows our BiophysicalNN to effectively learn the tradeoffs between negative and hydrophobic amino acids described above.

In conclusion, we advocate for a dual-pronged approach incorporating both biophysical NNs and deeper, more accurate NNs. The deep NN can provide the most accurate predictions, while the biophysical NNs can be used to understand the post-hoc interpretations of the deep NN. As NNs become a larger part of biological research, we hope this multi-pronged interpretation will provide another avenue to better understand biology.

## Acknowledgments

We thank James Galagan, Hunter Nisonoff, and James Bowden for very helpful discussions. We thank Tomas Rube and Ryan Emenecker for feedback on the manuscript and members of the Staller Lab for helpful discussions. C.J.L. was supported by NIH T32HG4725. P.A. was supported by UC Berkeley SURF L&S fellowship. G.H. was supported by a UC Berkeley Rose Hills summer scholarship. J.D. was supported by UC Berkeley CDSS. M.Z. was supported by NIH T32GM148378. A.L. was supported by UC Berkeley URAP. J.E.C.H. was supported by the UC Berkeley SEED Scholars Honors Program. M.V.S. was supported by NSF grant 2112057, and NIH grant R35GM150813. M.V.S. is a Chan Zuckerberg Biohub – San Francisco Investigator.

## Methods

### Dataset preparation

The dataset used for training and testing is from our previous screen^17^. The 40-amino acid sequences originate from homologs of the *S. cerevisiae* Gcn4 transcription factor, a stress-response TF that contains a highly studied acidic activation domain. The full length Gcn4 homolog sequences are broken into 40 amino acid tiles, separated by five amino acids. This tiling strategy resulted in 20731 designed 40 amino acid tiles. Sort-seq^23,26^ was performed to determine the GFP activation and mCherry abundance measurements for each tile (**Figure 1A**). In total, we had mCherry and GFP data for 17732 tiles. When tiles were present in both of the two replicates, we averaged the score from each replicate to get a single score for each tile.

Because of the tiling process, many sequences in our dataset were closely related, with some differing by as few as 5 amino acids. This overlap means that if our NN memorized the training data, it may still perform well on the test dataset, depending on how the sequences were split into train/test splits. To account for this potential bias, we split the data into train, validation and test sets with minimal overlap. We hierarchically clustered the full length ortholog sequences, split the sequences into three clusters, and then assigned tiles to each cluster based on the ortholog sequence that they originated from (**Figure S13A**). In this way, tiles from the same full length sequence would not be in both the training, validation and the test split. Additionally, we added all control sequences to the test set (**Figure S13A**). As a final test, we verified that there was minimal correlation between the similarity of sequences in the test set to the training set and the prediction accuracy (**Figure S13B**).

### Neural Network design

For all the NN variations, we used the same basic building block, illustrated in **Figure 1B**. This block takes as input the one-hot encoded activation domain sequence and predicts either the total TF abundance (mCherry fluorescence), the activation domain strength (GFP fluorescence) or an intermediate biophysical value (the equilibrium constant between states).

The NNs were created using PyTorch^60^. Additionally, code to load the data was modified from Faure and Lehner^11^. For the SimpleNN, it was straightforward to use a single block to predict either the GFP activation or the TF abundance.

To predict these measurements using the biophysical models, we first derived an equation relating the biophysical models to the experimental data. This required two assumptions (**Figure 3B, 4B**). First, the total mCherry fluorescence is equal to the amount of transcription factor in each state. Second, the total GFP fluorescence is related to the amount of transcription factor in the bound state through some function. We compared a linear function and a hill function and found that the hill function led to the best performance (**Figure S14**).

Finally, we assume that the states of the biophysical models are in equilibrium. Using these three assumptions, we derived an equation relating the equilibrium constants to the experimental measurements (**Figure S7**).

### Neural Network training

For each NN, we trained thirty identical models with different random seeds. We trained for 200 epochs with early stopping. We used a learning rate of 0.001, as that led to highest performance. For the SimpleNN, our loss function was the mean absolute error (MAE) between the predicted and actual values.

For the BiophysicalNN, we performed the training in two stages, because we were predicting two different values. First, we trained the abundance predictor. We froze all the weights not involved in predicting the abundance and used a loss function that only included the abundance values. This loss function also included a term to penalize negative linear weights (this helped in interpretability and did not have a noticeable impact on performance). Specifically, the loss function was:

**loss(predicted abundance, experimental abundance) = MAE(predicted abundance, experimental abundance) + (- sum of negative weights)**

After we had trained the abundance module, as defined by the validation loss no longer improving, we trained the activation domain strength prediction module. We froze the weights involved in predicting the abundance, and used a loss function that only included activation domain strength values. We noticed that the best performing NN often predicted negative equilibrium constants, which does not make sense from a biological perspective. Therefore, we also added a term to the loss function to penalize negative equilibrium constants. Similar to above, we also included a term in the loss function to penalize negative linear weights, which helped with interpretability. The loss function for this stage of training was:

**loss(predicted activation, experimental activation) = MAE(predicted activation, experimental activation) + 0.1*(- sum of negative weights) + 0.1*(- sum of negative equilibrium constants)**

### Details of how the NN layers work

In what follows, we describe how each of the layers of the network function. Although this part is likely unnecessary for readers experienced in ML, we include this section because an understanding of how each component of the network functions is essential to interpret the parameters of the network.

Each NN begins by encoding the input sequence with a one-hot encoding. A one-hot encoding transforms the 40 amino acid sequence into a (20×40) matrix where the columns correspond to sequence position and the rows correspond to amino acids. A one in the matrix indicates the specified amino acid is in the given position.

A convolutional filter is then directly applied to the encoding. The convolutional filter is a (20 x k) matrix where the rows correspond to the twenty possible amino acids and the columns correspond to position in the sequence. Starting from the beginning of the sequence, this filter matrix is element wise multiplied with the portion of the encoded sequence that it overlaps with. If the filter is (20×5), it would be multiplied with the first five columns in the one hot encoding. The result of this multiplication is summed and a bias term is added, resulting in a single value. The filter then slides one position over (i.e. starts at the second amino acid position in the one hot encoding) and repeats the procedure, adding the same bias term. The filter continues until it reaches the end of the sequence. This results in a 40 - k + 1 length vector. In summary, the convolutional filter is a method of scanning the entire sequence for specific patterns/motifs/important residues without regard for where in the sequence these residues appear.

The Parametric ReLU activation function transforms each value in the 40 - k + 1 vector that resulted from the convolution. It is a variant of the ReLU activation function where positive values remain unchanged while negative values are shrunk by some scaler. Activation functions allow NNs to learn non-linear relationships between sequence and biophysical value. After this activation function is applied, the resulting vector is multiplied by weights with some bias added. These linear weights serve to weigh patterns captured by the convolution filter by their location in the sequence, capturing positional effects. Finally, a second Parametric ReLU function is applied.

### ADhunter Implementation

ADhunter is a residual neural network implemented in PyTorch^60^ and trained via PyTorch Lightning^61^. Its architecture consists of an embedding layer, a convolutional layer, three residual blocks (batch norm layer, convolutional layer, another batch norm layer), followed by a max pool layer and a dense layer. It is trained using the same train/validation/test splits as our SimpleNNs and BiophysicalNNs with mean squared error loss and early stopping. We retrained ADhunter on each dataset (GFP, mCherry).

### Comparison with other NNs

One difficulty in directly comparing various activation domain predictors is that each predictor takes a different input size: TADA takes 40aa long inputs, PADDLE requires a 53aa long input and ADPred takes any input size but produces a predicted value for each residue instead of a single score for the entire sequence. We ran TADA and ADPred on the 40aa tiles in the test set. To get a single value for ADPred, we averaged the predictions for the 40aa tile. To run PADDLE, we also included the 13 amino acids in the synthetic TF N-terminal to the 40aa tile to create a 53aa tile.

### Deep NN interpretations

DeepLIFT^9^ quantifies the effect of each input feature on the final prediction. Specifically, it calculates how much each input feature contributes to the difference in scores between the model evaluated with the input and some baseline. For our analysis, we used a null baseline where our one-hot encoded matrix is simply a matrix of zeros. We performed this DeepLIFT analysis on all active sequences in the Gcn4 dataset^17^ and averaged the result to get a summary of global behavior. More details and the code used to perform the analysis is available on github.

To interpret PADDLE^25^, we took five Gcn4 homologs and we performed all possible single amino acid mutations. To investigate the effect of position within the 53 amino acid input, we centered the mutated amino acid at every possible position within the 53mer. This lead to a total of 235,320 mutated tiles. After running PADDLE on all of these tiles, we calculated the change in activation domain strength between the wildtype and mutant tiles. We then grouped the tiles by mutation position within the tile and the mutated residue identity, calculating the average change in predicted strength for each group.

### FINCHES predictions

We used FINCHES^40^ to predict the interaction between the ABD1 domain of Med15 and all the tiles in our dataset using the folded domain surface interactions method. Specifically, FINCHES identifies solvent accessible patches of the structured protein and predicts interactions between these patches and the IDR. We looked at only the attractive score between the coactivator and activation domain, as the attractive score showed the best correlation across all comparisons.

## Data and Code availability

- The raw sequencing data has been deposited at NIH SRA Accession #PRJNA1186961: http://www.ncbi.nlm.nih.gov/bioproject/1186961
- All code and processed used in this analysis are deposited at GitHub: https://github.com/staller-lab/NN_interpretability_for_AD_prediction and Zenodo: 10.5281/zenodo.17102617

## Supplemental Figures

**Figure S1:**
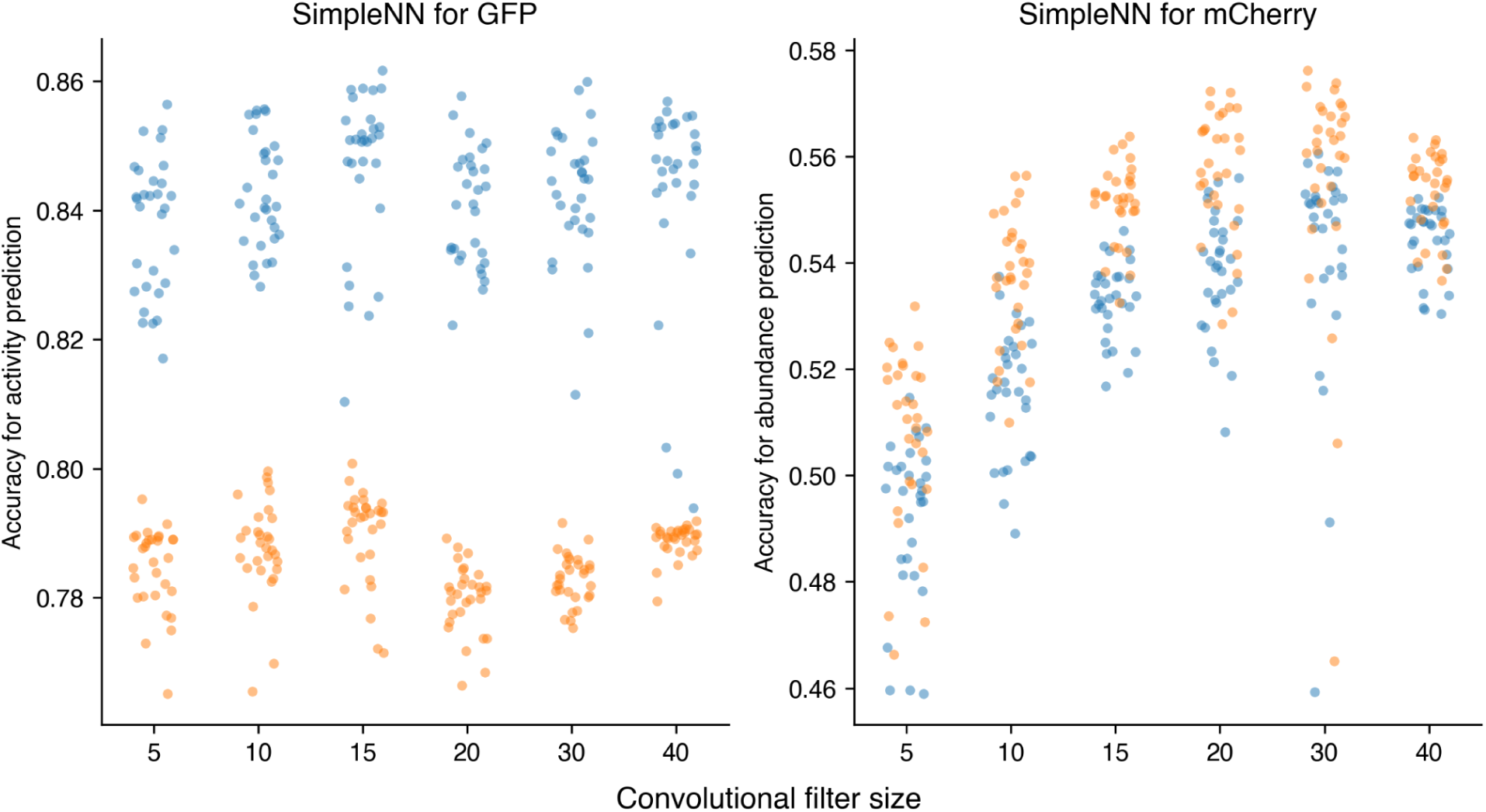
Random initialization of the SimpleNN has a large impact on the performance. The orange points are showing the spearman correlations and the blue are showing the Pearson correlations. For each convolutional filter size, the same NN architecture was trained thirty times, varying only the random seeds. All filter sizes perform similarly for the activation domain strength prediction task. For abundance prediction, the size 20 and 30 kernels seem to perform slightly better, although random initialization still has a large impact of final prediction.

**Figure S2:**
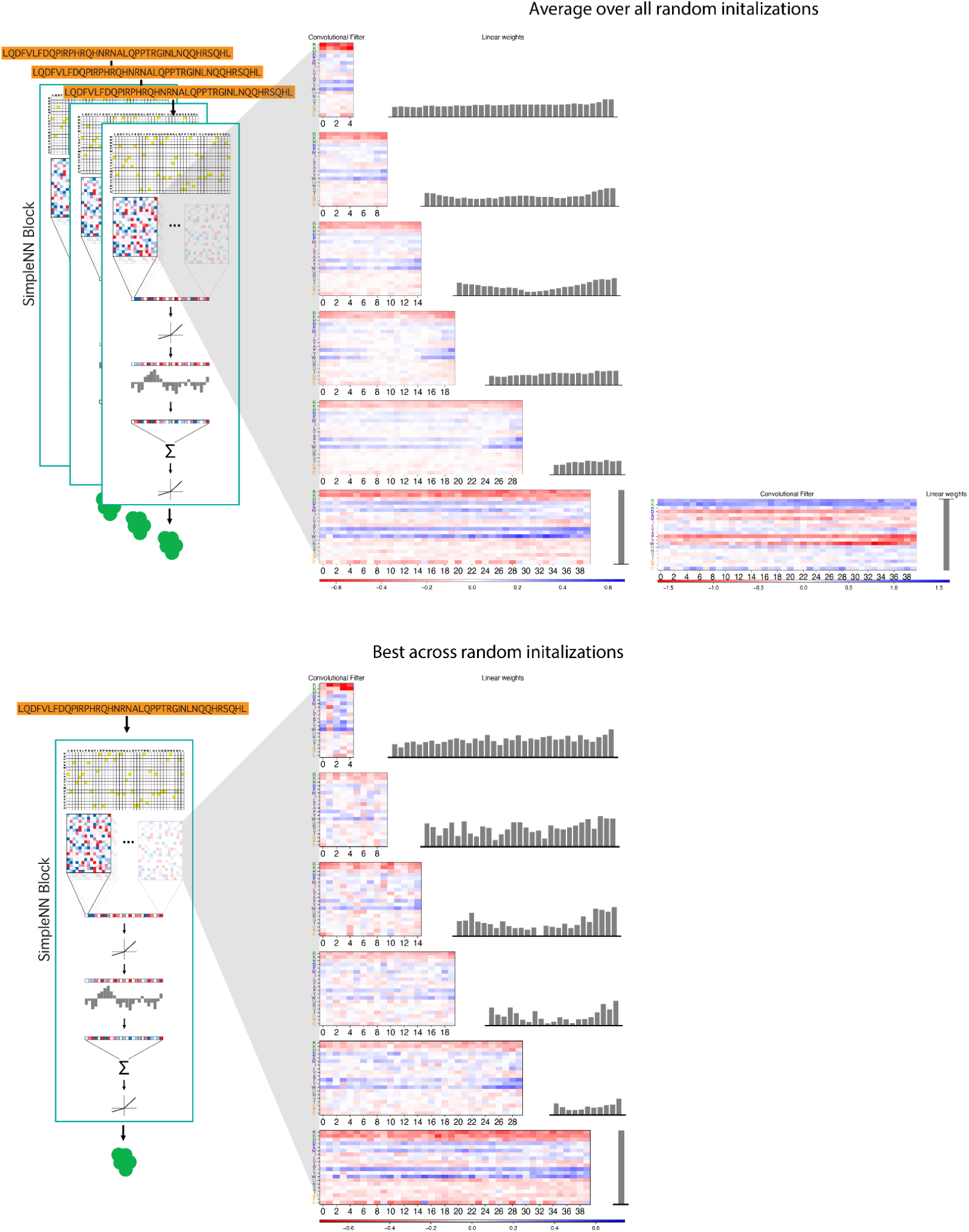
Parameters of the SimpleNN for GFP across all random initializations. Row 1: For each convolutional filter size (5-40) for the SimpleNN, we average the convolutional filters and weights of the thirty NNs with different random seeds. The NNs with negative and positive linear weights were averaged separately (column 1 and column 2, respectively) as we anticipate the convolutional filters to learn opposite weights. Row 2: The parameters of the best performing NNs for each filter size.

**Figure S3:**
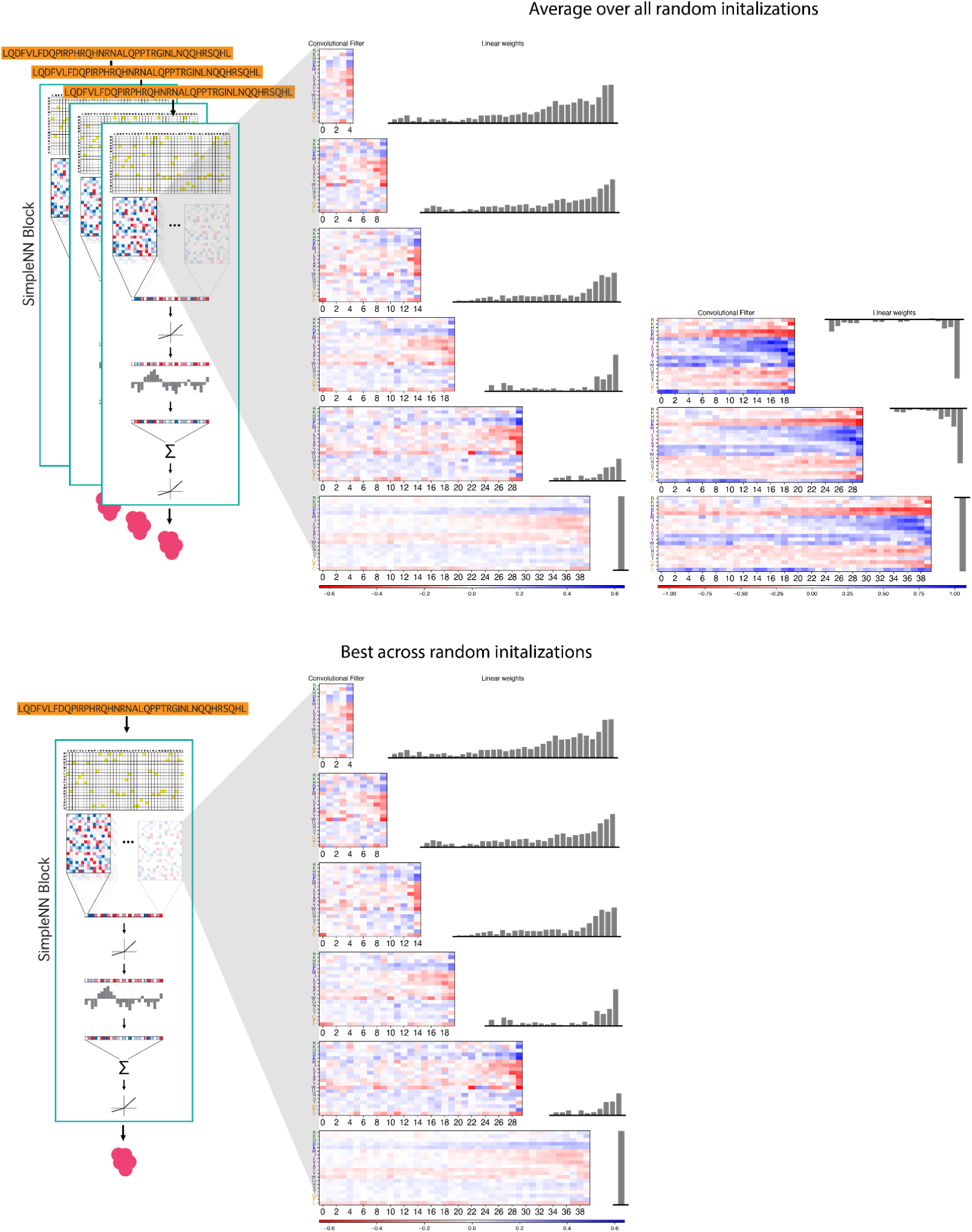
Parameters of the SimpleNN-abund across all random initializations. Row 1: For each convolutional filter size (5-40) for the SimpleNN-abund, we average the convolutional filters and weights of the thirty NNs with different random seeds. The NNs with negative and positive linear weights were averaged separately (column 1 and column 2, respectively) as we anticipate the convolutional filters to learn opposite weights. Row 2: The parameters of the best performing NNs for each filter size.

**Figure S4:**
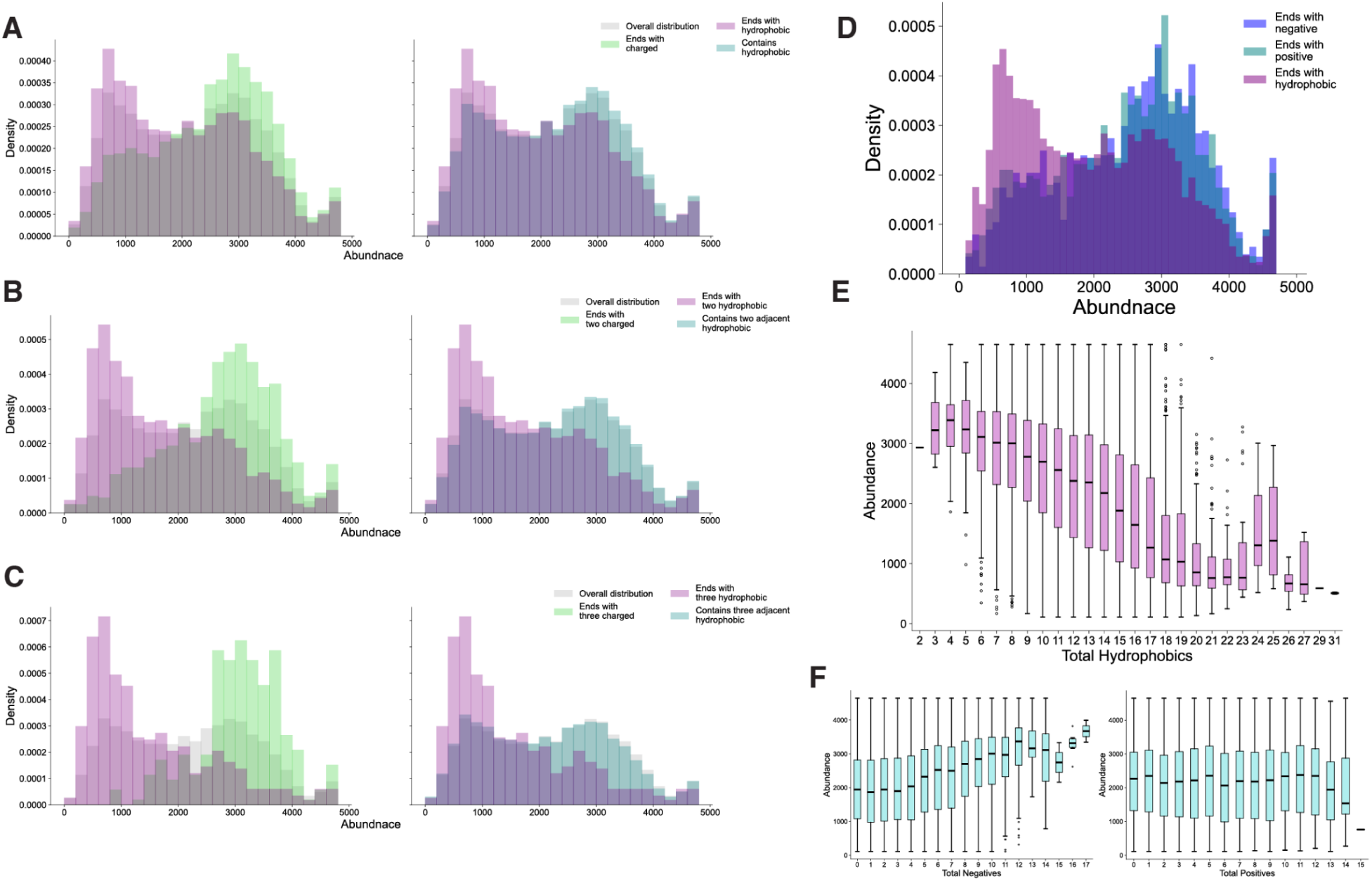
Transcription factor abundance is influenced by overall hydrophobic content and hydrophobic amino acids at the C-termini. In all plots, hydrophobic amino acids are M, I, L, V, A, F, Y, W. Positive amino acids are R and K. Negative amino acids are D and E. Charged amino acids are positive and negative amino acids. **A)** The abundance distributions for transcription factors ending in one hydrophobic amino acid versus one charged amino acid. The second plot compares the abundance distributions for ending in one hydrophobic amino acid versus containing one hydrophobic amino acid anywhere in the sequence. **B-C)** The same as **A** but for two and three charged/hydrophobic respectively. The second column compares two/three adjacent hydrophobic amino acids anywhere in the sequence to two/three at the end of the sequence. **D)** The abundance distribution for sequences ending in a hydrophobic amino acid versus negative amino acids versus positive amino acids. Both positive and negative amino acids increase abundance. **E)** Box plot showing how abundance distributions change based on the number of overall hydrophobic amino acids in the sequence. **F)** Boxplots showing how abundance distributions changed based on the overall number of negative and positive amino acids in the sequence.

**Figure S5:**
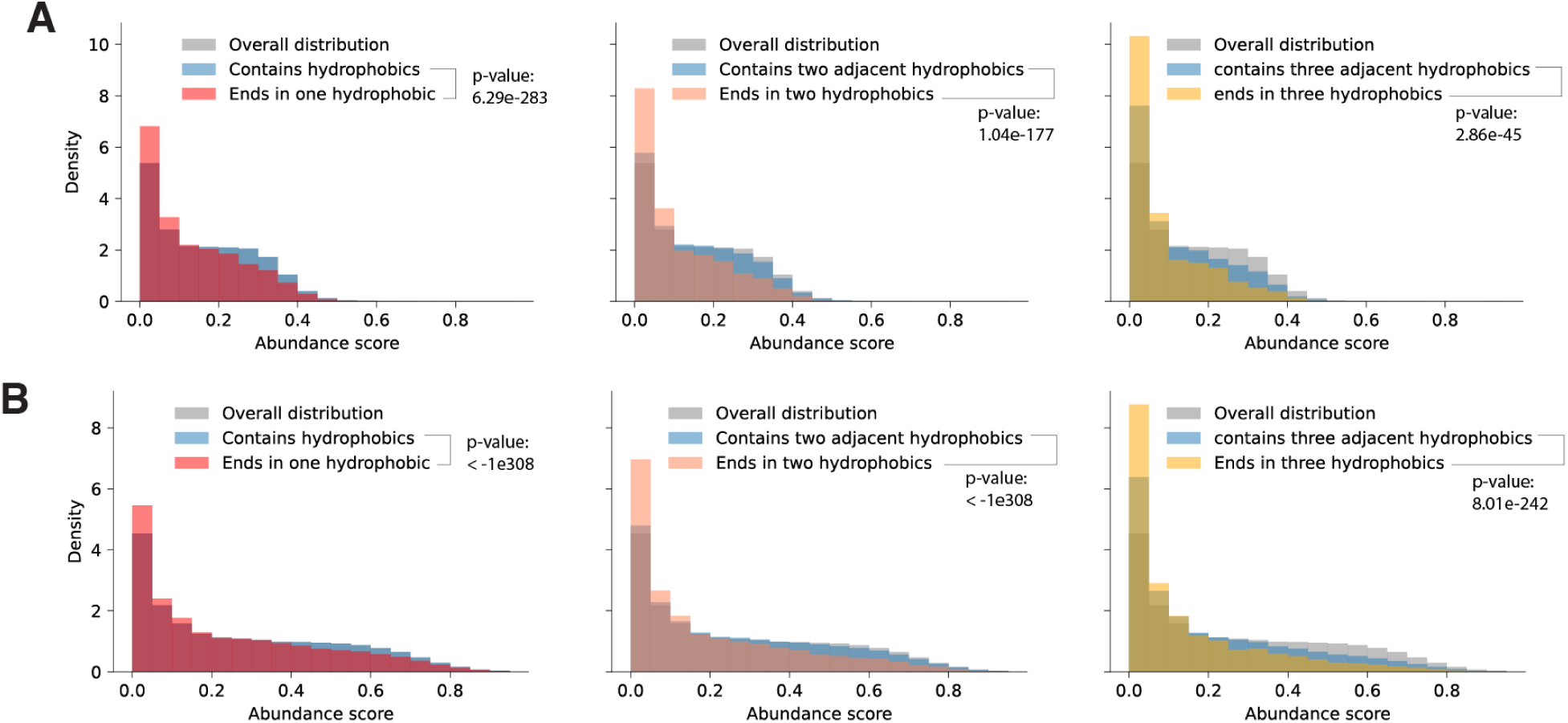
Other degron screens show similar C-terminal hydrophobic effects. **A.** Distribution of abundances from Voutsinos et al.^36^ indicates that proteins with C-terminal hydrophobic amino acids have decreased abundance compared to other proteins. *p*-values are from a one-tailed Mann-Whitney U test. **B.** Transcription factor degron screen data from Larsen et al.^35^ also shows similar C-terminal effects. *p*-values are from a one-tailed Mann-Whitney U test.

**Figure S6:**
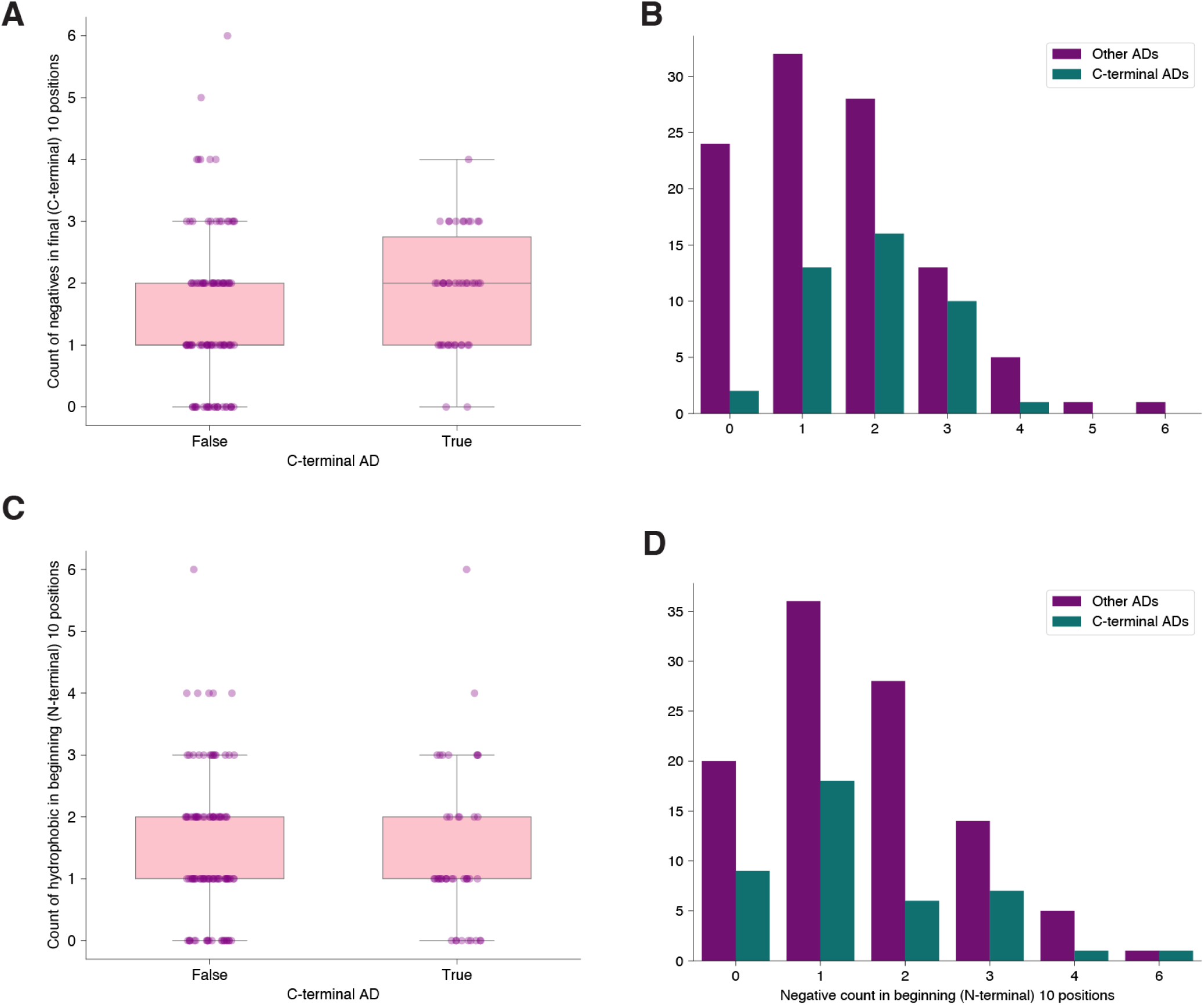
C-terminal activation domains are enriched in C-terminal negative charge. C-terminal activation domains were defined as activation domains that were within 3 amino acids of the end of the sequence. **A)** The non-C-terminal activation domain set had significantly less negative charge in the last 10 amino acids than the C-terminal activation domain set (*p*-value = 0.015, Wilcoxon rank sum test, one-tailed). **B)** Histogram showing the same data as in A. **C)** There was no significant difference in the number of negatives in the ten most N-terminal most amino acids between the C-terminal vs. non-C-terminal activation domains (*p*-value = 0.505, Wilcoxon rank sum test). **D**) The same data in C plotted as a histogram.

**Figure S7:**
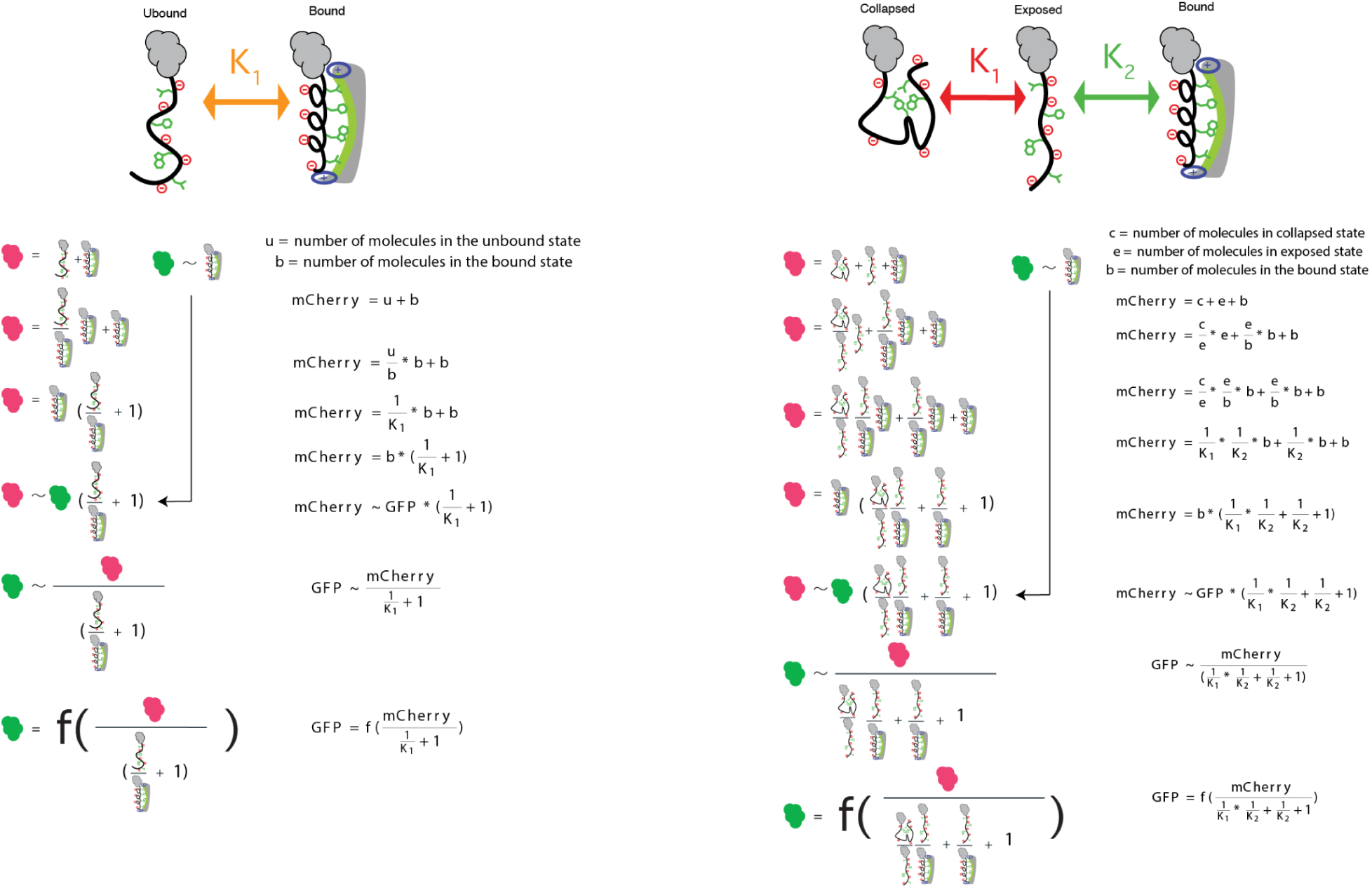
Derivation of the binding equations used in the BiophysicalNNs. These equations were derived to relate the biophysical models to the amount of mCherry and GFP observed. The same derivation is show using math and cartoons. Left: Two-state biophysical model. Right: Three-state biophysical model.

**Figure S8:**
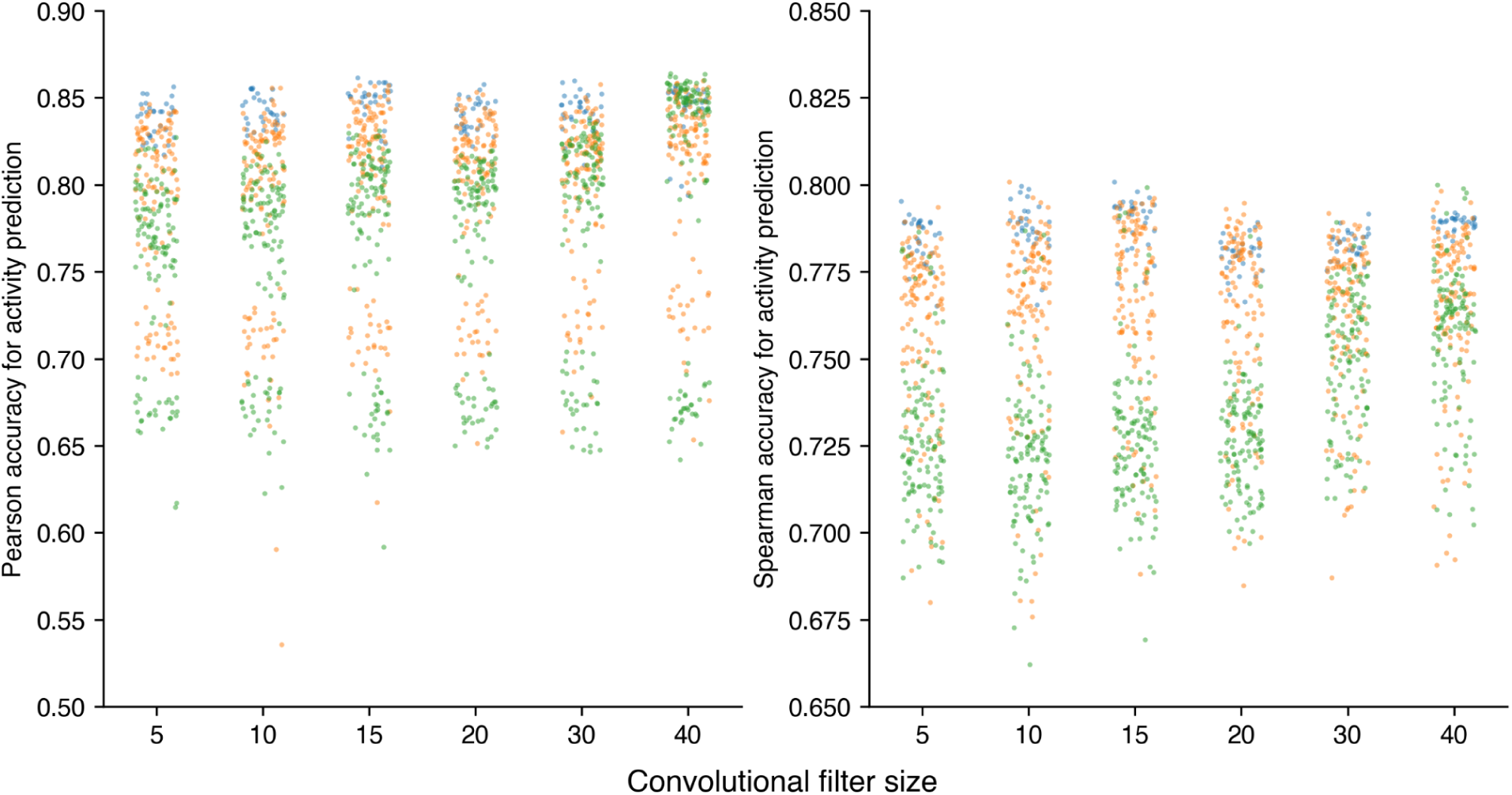
The three-state BiophysicalNN performs better than the two-state BiophysicalNN at smaller filter sizes. Blue points show the SimpleNN, orange points show the three-state BiophysicalNN and green points show the two-state BiophysicalNN. For each filter size, the two and three-state BiophysicalNNs were trained 180 times, varying only the different random seeds and the abundance filter size. The SimpleNN was trained 30 times per filter size. The Pearson and spearman correlations for predicting the activation domain strength are shown on the left and right, respectively. Note that the two-state BiophysicalNN only performs well at larger convolutional filter sizes, likely because only these filter sizes are able to account for positional effects of certain amino acids.

**Figure S9:**
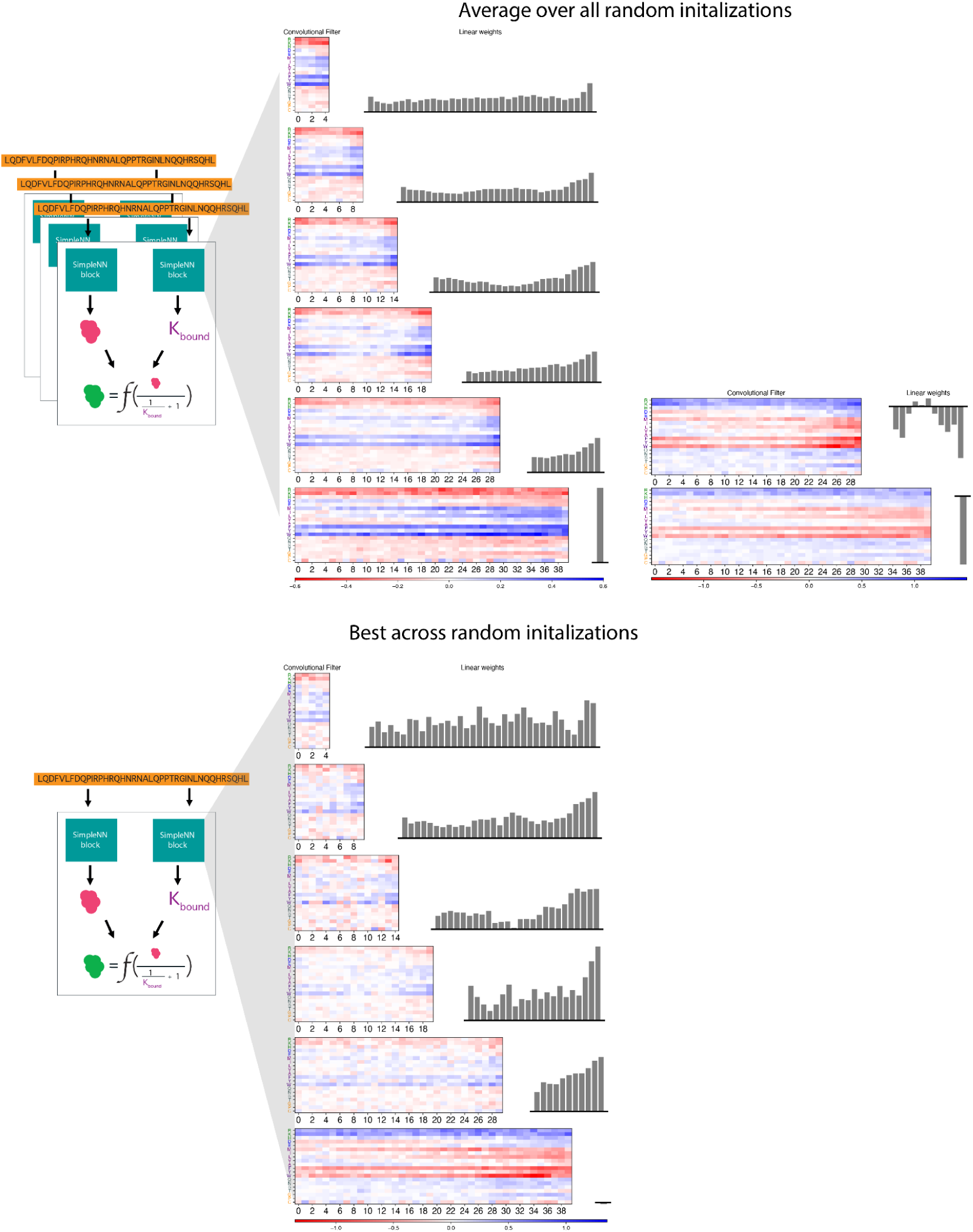
Parameters of the two-stateBiophysicalNN across all random initializations. Convolution filters and linear weights for the SimpleNN block predicting the equilibrium constant of the two-state biophysical model. First row: For each convolutional filter size (5-40) for the two-state BiophysicalNN, we average the convolutional filters and weights of the 180 NNs with different random seeds and abundance filter sizes. The NNs with negative and positive linear weights were averaged separately (column 1 and column 2, respectively) as we anticipate the convolutional filters to learn opposite weights. Second row: The parameters of the best performing NNs for each filter size.

**Figure S10:**
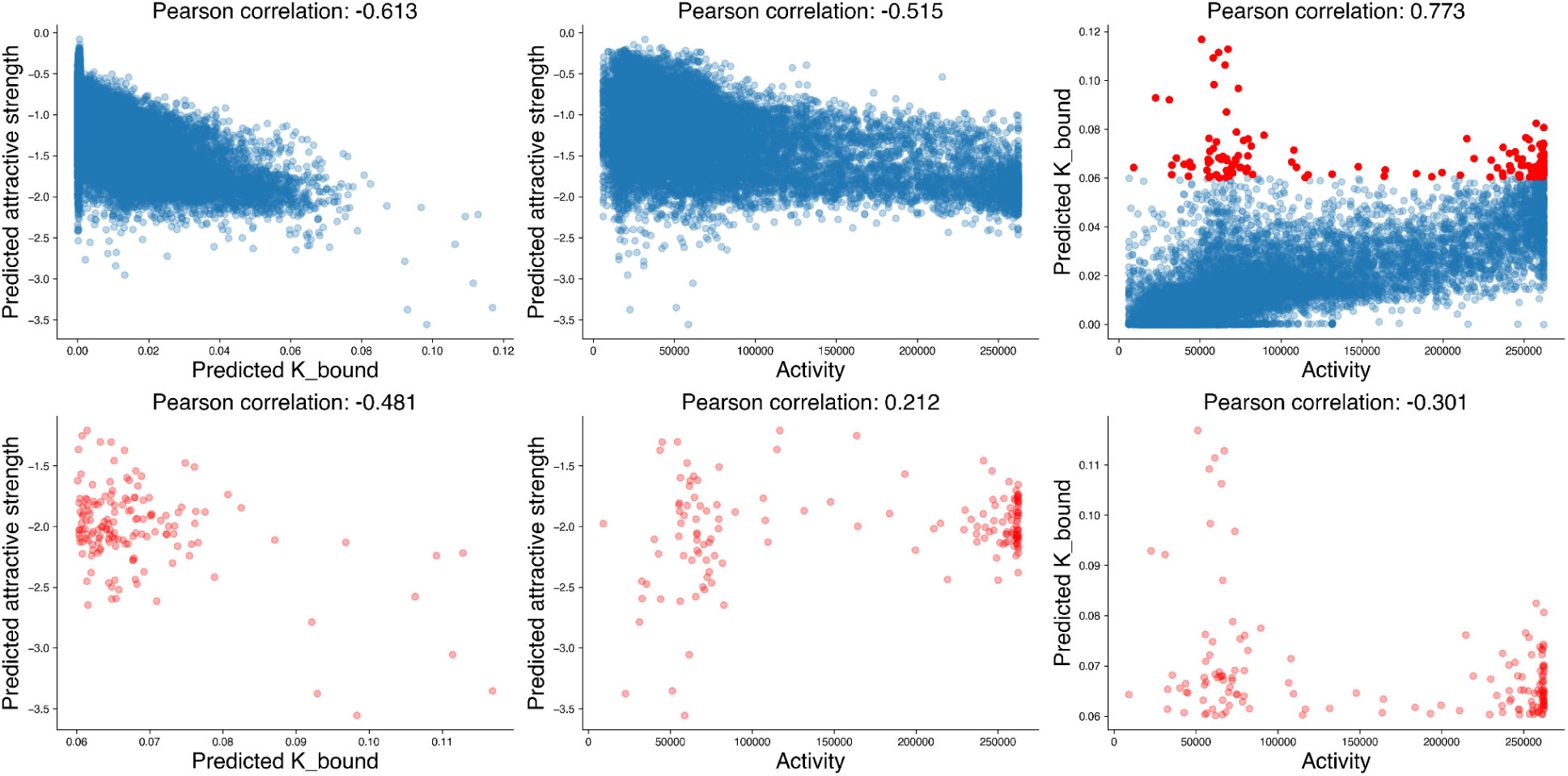
Equilibrium constant between unbound and bound states is correlated with FINCHES^40^ predicted interaction strength more than with activation domain strength. Equilibrium constants are from the two-state BiophysicalNN. A more negative attractive strength indicates a stronger predicted interaction. The Calvados force field^62^ was used. The bottom row shows a subset of sequences with high equilibrium constants (K > 0.06). In these sequences, only the correlation between FINCHES score and the equilibrium constant is maintained.

**Figure S11:**
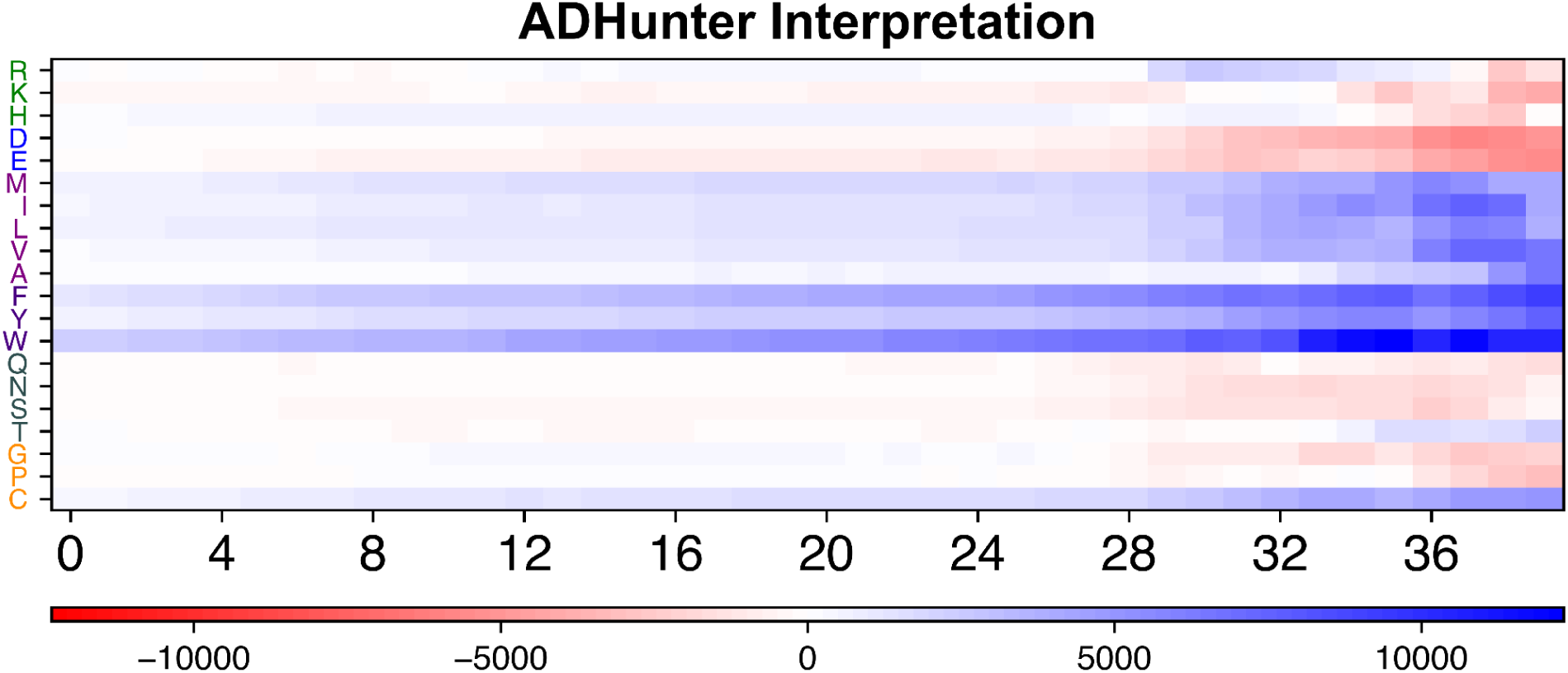
Post-hoc interpretation of ADhunter-ratio. This interpretation was performed using the method used to interpret PADDLE, by quantifying the effect of all possible mutations in the same five sequences. This analysis captures similar trends to the DeepLIFT analysis of ADhunter, such as the detrimental impact of C-terminal negative amino acids and the strong beneficial impact C-terminal hydrophobic amino acids. Interestingly, it does not capture the bifunctional role of negative amino acids seen in the ADhunter DeepLIFT. This discrepancy may be due to the minimal number of sequences used in this analysis compared to DeepLIFT. As this analysis is very computationally intensive, it was not feasible to run it on a similar number of sequences, but we are encouraged that they are both capturing similar C-terminal effects.

**Figure S12:**
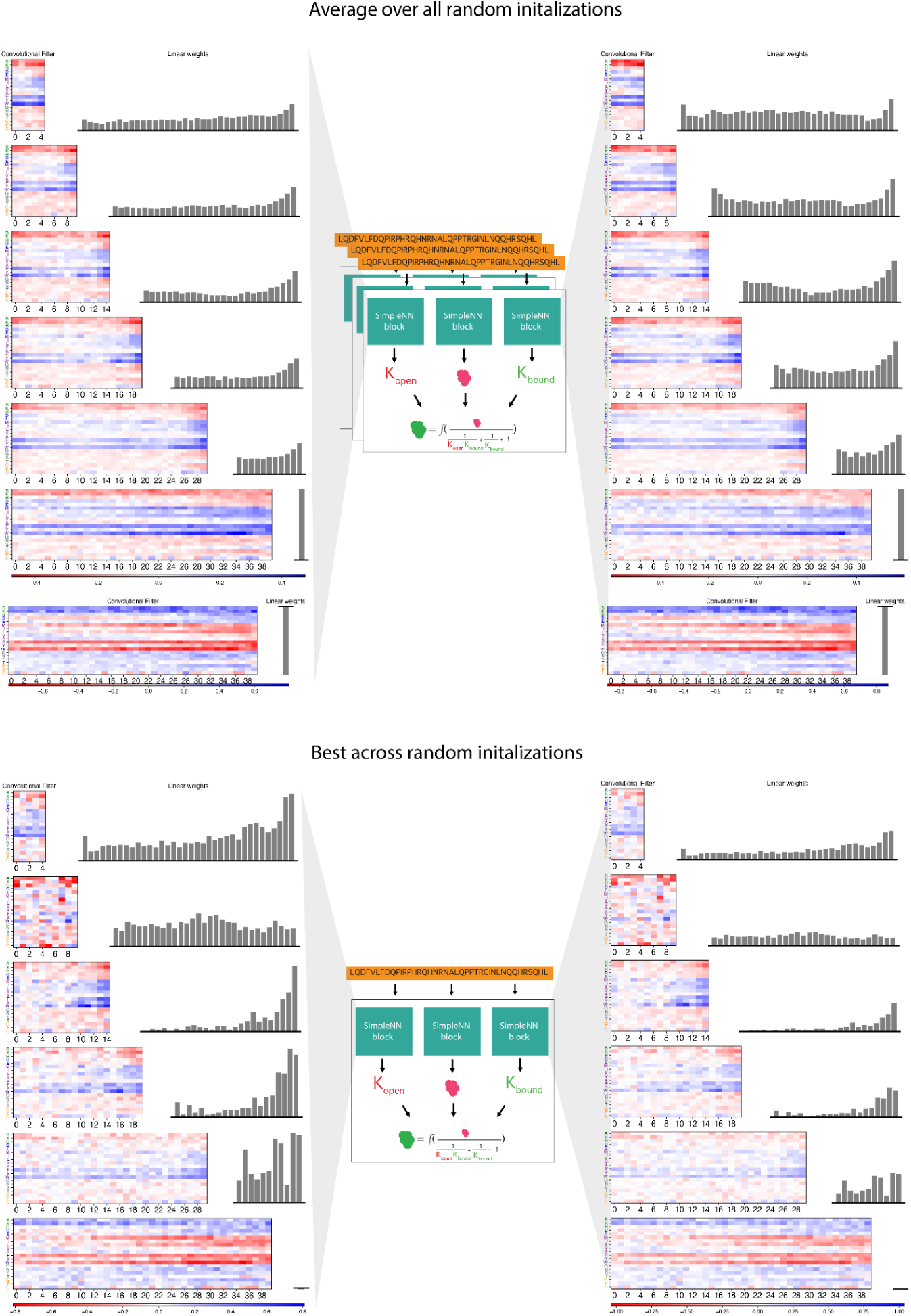
Parameters of the three-stateBiophysicalNN across all random initializations. Row 1: For each convolutional filter size (5-40) for the three-state BiophysicalNN, we average the convolutional filters and weights of the 180 NNs with different random seeds and abundance filter sizes. The NNs with negative and positive linear weights are shown separately. Row 2: The parameters of the best performing NNs for each filter size.

**Figure S13:**
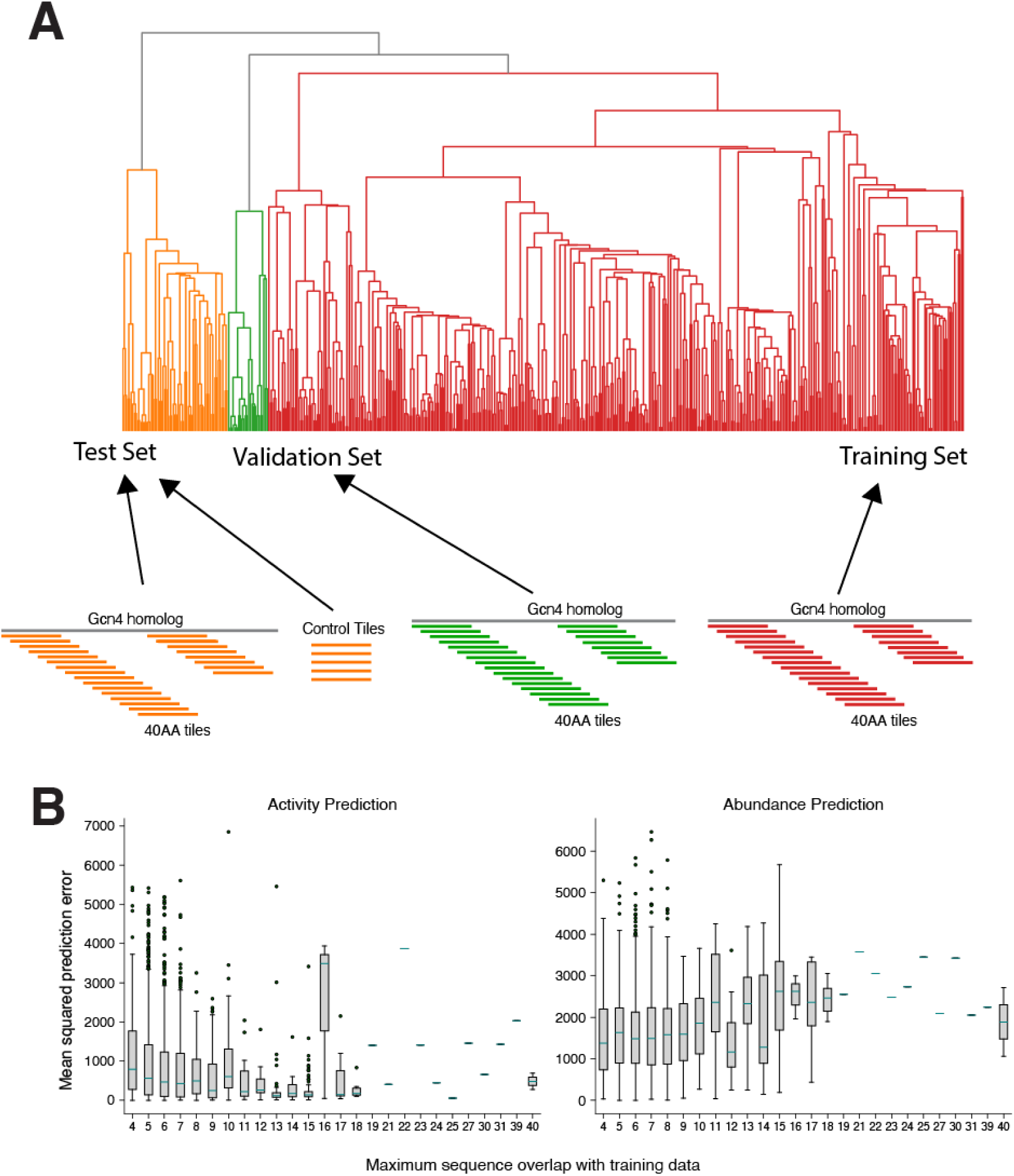
Scheme for ensuring minimal overlap between the test, validation and training data. **A)** The full length transcription factors were hierarchically clustered based on their amino acid sequences. We split the three top-most clusters into the training, validation and test datasets. Tiles were then assigned to each set based on the full length TF that they originated from. Control tiles were added to the test dataset. **B)** Predictive performance on the test dataset is minimally affected by similarity to the training dataset.

**Figure S14:**
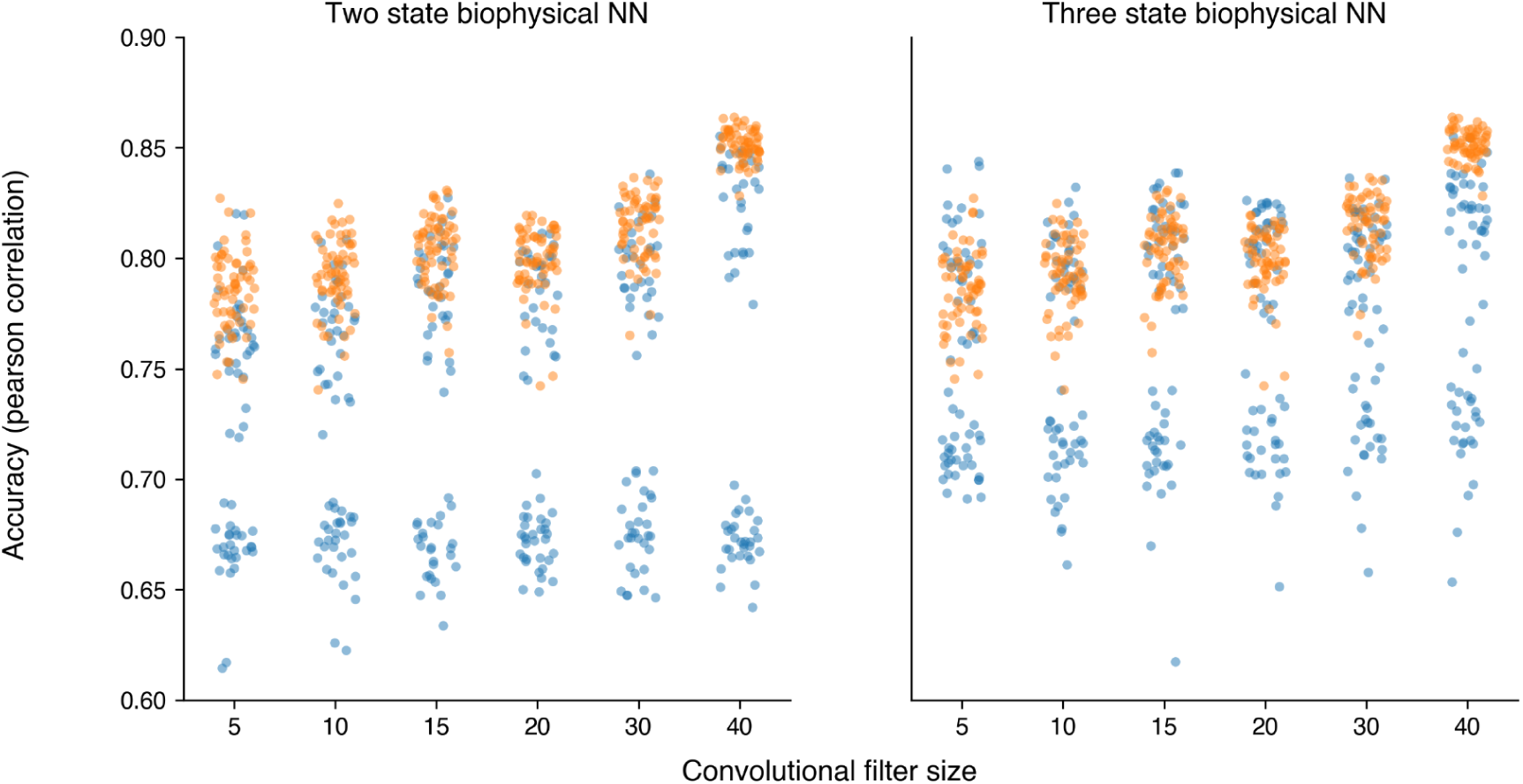
Hill function outperformed the linear function for transforming the amount of bound transcription factor to the amount of reporter gene expression. For each activation function (hill and linear) and convolutional filter size, thirty NNs were trained varying only the random initialization. Blue points indicate performance with the linear function and orange points indicate performance with the hill function. The Pearson correlation for predicting the GFP expression is shown.

## Supplemental Tables

**Table S1:** The NNs generalize similarly to deep NNs. Here, we test NN performance on two recent datasets measuring the GFP expression of activation domains. These datasets are of lower quality than the dataset used in this study, so all NNs suffer a performance hit. Hummel data from Hummel et al.27 Because they measured GFP for 53aa long tiles, for TADA, ADhunter, the SimpleNN, and both BiophysicalNNs (which all take 40aa inputs), we tiled the 53aa input, ran the predictors on each tile, and then averaged the predictions. PADI data from GFP data from Morffy et al.28 PADI data measured 40aa tiles. Since PADDLE takes a 53aa input, we predicted on the most C-terminal 53aa in the synthetic TF. ADPred predicts a score for each position, so to get a single value for comparison, we took the maximum predicted value, as this showed the best performance.

| Dataset | NN | Pearson | Spearman | PRC-AUC | ROC-AUC |
| --- | --- | --- | --- | --- | --- |
| Hummel | Simple | 0.290 | 0.268 | 0.718 | 0.599 |
| Hummel | Two-state | 0.326 | 0.302 | 0.731 | 0.613 |
| Hummel | Three-state | 0.312 | 0.291 | 0.725 | 0.606 |
| Hummel | ADhunter-GFP | 0.362 | 0.335 | 0.744 | 0.624 |
| Hummel | PADDLE | 0.340 | 0.339 | 0.756 | 0.639 |
| Hummel | TADA | N/A | N/A | 0.738 | 0.609 |
| Hummel | ADPred | N/A | N/A | 0.705 | 0.579 |
| PADI | Simple | 0.461 | 0.356 | 0.421 | 0.813 |
| PADI | Two-state | 0.473 | 0.333 | 0.443 | 0.808 |
| PADI | Three-state | 0.485 | 0.358 | 0.455 | 0.818 |
| PADI | ADhunter-GFP | 0.520 | 0.386 | 0.496 | 0.831 |
| PADI | PADDLE | 0.520 | 0.386 | 0.514 | 0.835 |
| PADI | TADA | N/A | N/A | 0.555 | 0.859 |
| PADI | ADPred | N/A | N/A | 0.461 | 0.828 |

